# Tumour mitochondrial DNA mutations drive aerobic glycolysis to enhance checkpoint blockade

**DOI:** 10.1101/2023.03.21.533091

**Authors:** Mahnoor Mahmood, Eric Minwei Liu, Amy L. Shergold, Elisabetta Tolla, Jacqueline Tait-Mulder, Alejandro Huerta Uribe, Engy Shokry, Alex L. Young, Sergio Lilla, Minsoo Kim, Tricia Park, J.L. Manchon, Crístina Rodríguez-Antona, Rowan C. Walters, Roger J. Springett, James N. Blaza, Sara Zanivan, David Sumpton, Edward W. Roberts, Ed Reznik, Payam A. Gammage

**Affiliations:** Cancer Research UK Beatson Institute, Glasgow, UK; Computational Oncology Service, Memorial Sloan Kettering Cancer Center, New York, NY, USA; Centro Nacional de Investigaciones Oncológicas(CNIO), Madrid, Spain; Centro de Investigación Biomédica en Red de Enfermedades Raras CIBERER, Madrid, Spain; Structural Biology Laboratory and York Biomedical Research Institute, Department of Chemistry, The University of York, York, UK; School of Cancer Sciences, University of Glasgow, UK; Marie-Josée and Henry R. Kravis Center for Molecular Oncology, Memorial Sloan Kettering Cancer Center, New York, NY, USA; Urology Service, Memorial Sloan Kettering Cancer Center, New York, NY, USA

## Abstract

The mitochondrial genome encodes essential machinery for respiration and metabolic homeostasis but is paradoxically among the most common targets of somatic mutation in the cancer genome, with truncating mutations in respiratory complex I genes being most over-represented^1^. While mitochondrial DNA (mtDNA) mutations have been associated with both improved and worsened prognoses in several tumour lineages^1–,3^, whether these mutations are drivers or exert any functional effect on tumour biology remains controversial. Here we discovered that complex I-encoding mtDNA mutations are sufficient to remodel the tumour immune landscape and therapeutic resistance to immune checkpoint blockade. Using mtDNA base editing technology^4^ we engineered recurrent truncating mutations in the mtDNA-encoded complex I gene, *Mt-Nd5*, into murine models of melanoma. Mechanistically, these mutations promoted utilisation of pyruvate as a terminal electron acceptor and increased glycolytic flux without major effects on oxygen consumption, driven by an over-reduced NAD pool and NADH shuttling between GAPDH and MDH1, mediating a Warburg-like metabolic shift. In turn, without modifying tumour growth, this altered cancer cell-intrinsic metabolism reshaped the tumour microenvironment in both mice and humans, promoting an anti-tumour immune response characterised by loss of resident neutrophils. This subsequently sensitised tumours bearing high mtDNA mutant heteroplasmy to immune checkpoint blockade, with phenocopy of key metabolic changes being sufficient to mediate this effect. Strikingly, patient lesions bearing >50% mtDNA mutation heteroplasmy also demonstrated a >2.5-fold improved response rate to checkpoint inhibitor blockade. Taken together these data nominate mtDNA mutations as functional regulators of cancer metabolism and tumour biology, with potential for therapeutic exploitation and treatment stratification.

## Main

It has been known for several decades that >50% of cancers bear somatic mutations of mtDNA^5^. The impact of mtDNA mutations in the germline, the most common cause of inherited metabolic disease in humans^6^, is well-established. However, the biological and clinical relevance of mtDNA mutations in cancer remains contentious^5^. Recent efforts have yielded evidence for recurrence and selection of mtDNA mutations in cancer^1^, however the majority of variants observed somatically have not been detected in human disease or studied in the germline, thus requiring further study^1, 7^.

Hotspot truncating mutations in mitochondrial complex I genes are a common feature of several cancers, with truncating mutations in complex I (*MT-ND5* in particular) being over-represented compared with mutations in genes encoding respiratory complexes III, IV and V^1^. As complex I is a major site of NADH oxidation^8^ we reasoned that the proximal impact of complex I truncating mutations would be loss of NADH : ubiquinone oxidoreductase activity, resulting in redox imbalance with broad downstream impacts on cell metabolism. To test this hypothesis we designed mitochondria-targeted base editors^4^ to induce premature stop codons at tryptophan (TGA) codons within mouse *mt-Nd5*, analogous to hotspot mutations found in the human *MT-ND5* gene in tumours^1^ (**Figure 1A-C**).

**Figure 1.**
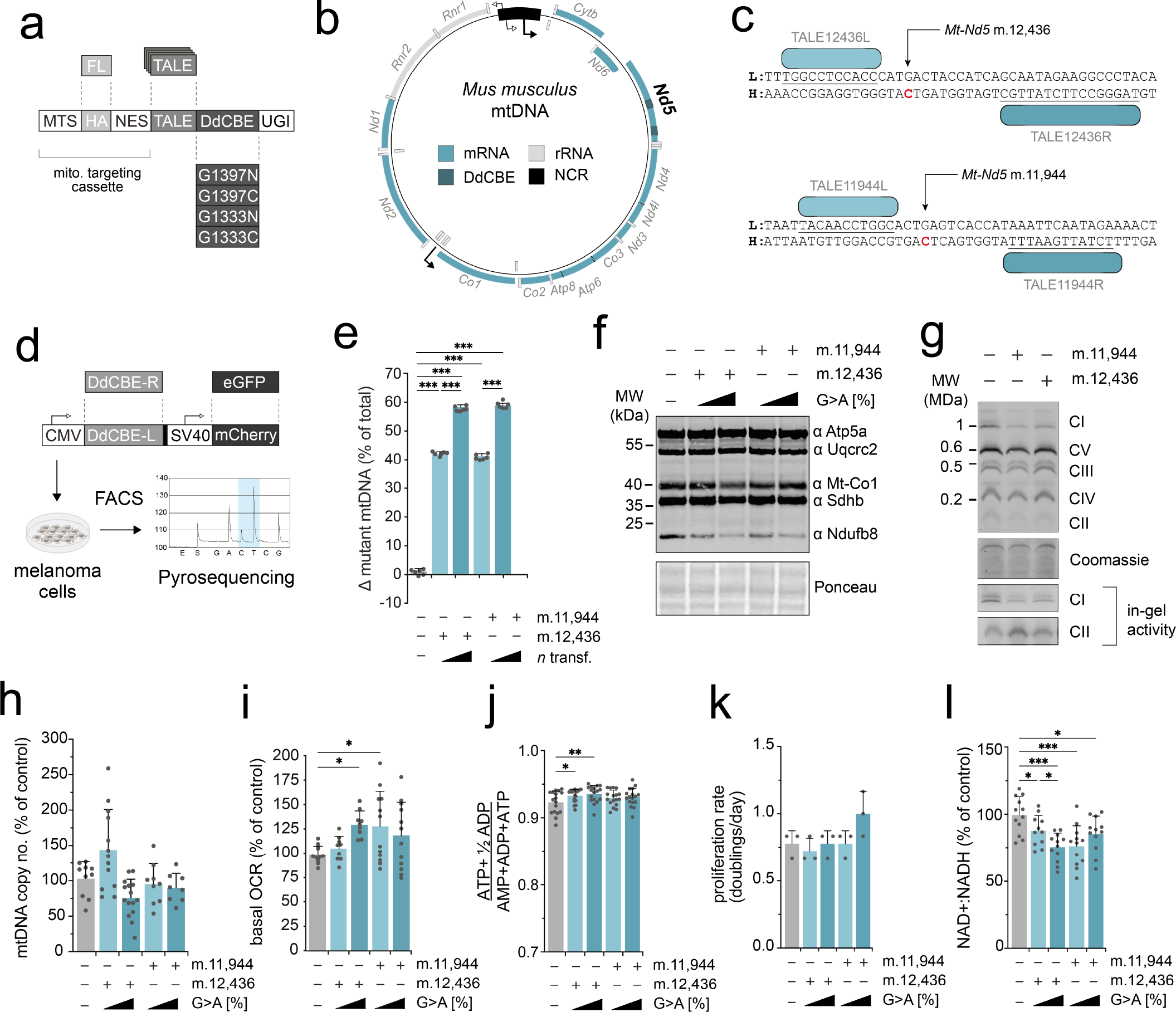
Mitochondrial base editing to produce isogenic cell lines bearing independent truncating mutation heteroplasmies in *mt-Nd5*. **A** Schematic of TALE- DdCBE design employed. TALEs were incorporated into a backbone containing a mitochondria-targeting cassette, split-half DdCBE and uracil glycosylase inhibitor (UGI). **B** Schematic of the murine mtDNA. Targeted sites within *mt-Nd5* are indicated. **C** TALE-DdCBE pairs used to induce a G>A mutation at mt.12,436 and mt.11,944. **D** Workflow used to produce *mt-Nd5* mutant isogenic cell lines. **E** Heteroplasmy measurements of cells generated in D (n = 6 separate wells were sampled). **F** Immunoblot of indicative respiratory chain subunits. Representative result is shown. **G** Assembled complex I abundance and in-gel activity of complexes I and II. Representative result is shown. **H** mtDNA copy number (n= 9 separate wells were sampled). **I** Basal oxygen consumption rate (OCR) (n = 9-12 measurements (12 wells per measurement) were made). **J** Energy (adenylate) charge state (n = 17 separate wells were sampled). **K** Proliferation rate of cell lines in permissive growth media. (n = 12 separate wells were measured in three batches) **L** NAD+:NADH ratio (n= 11-12 separate wells were measured). All P-values were determined using a one-way ANOVA test with (E, H-I, K) Sidak multiple comparisons test or (J,L) Fisher’s LSD Test. Error bars indicate SD. Measure of centrality is mean.

TALE-DdCBE G1397/G1333 candidates, bearing nuclear export signals, targeting m.12,436G>A and m.11,944G>A sites were synthesised and screened in mouse B78-D14 amelanotic melanoma cells (B.16 derivative, Cdkn2a null)^9^ to identify efficient pairs (**Figure 1D**). Expression of functional pairs (**Extended Data Figure 1A**) resulted in isogenic cell populations bearing ∼40% or ∼60% mutation heteroplasmy of m.12,436 G>A or m.11,944 G>A truncating mutations following either a single transfection or four consecutive transfections (referred to as m.12,436^40%^, m.12,436^60%^, m.11,944^40%^ and m.11,944^60%^ respectively) (**Figure 1E**) with limited off-target mutation (**Extended Data Figure 1B**). The resulting stable, isogenic cell lines demonstrated a heteroplasmy-dependent decrease in expression of complex I subunit Ndufb8 without impact on other respiratory chain components (**Figure 1F**). This was supported by Tandem Mass Tagging (TMT)-based mass spectrometry proteomics (**Extended Data Figure 2**) and blue native PAGE analysis of the m.12,436^60%^ and m.11,944^60%^ cell lines (**Figure 1G**), confirming that individual complex I subunit abundance, in addition to the proportion of fully assembled complex I, is decreased without substantial impact on other components of the OXPHOS system. In-gel activity assays of complex I and complex II activity further support this finding (**Figure 1G**). mtDNA copy number was not impacted by mutation incidence or heteroplasmy level (**Figure 1H**) and *mt-Nd5* transcript level was unchanged in m.12,436^60%^ and m.11,944^60%^ mutant cells compared with controls, consistent with lack of nonsense-mediated decay in mammalian mitochondria (**Extended Data Figure 3A**). Interestingly, none of the heteroplasmic cells exhibited significant decreases in oxygen consumption (**Figure 1I**), adenylate energy charge state (**Figure 1J**) or cell proliferation (**Figure 1K**). However, a ∼10mV decrease in the electrical component of the mitochondrial proton motive force , Δ^Ψ^, coupled to a commensurate trend towards ∼10mV increases in the chemical component , ΔpH, resulting in an unchanged total protonmotive force, Δ*P*, was detected (**Extended Data Figure 3B**). The NAD+ : NADH ratio was significantly impacted in mutant cells (**Figure 1L**), which was also reflected in reduced : oxidised glutathione (GSH : GSSG) ratios (**Extended Data Figure 3C**). The effect on cellular redox poise was further determined in m.12,436^60%^ and m.11,944^60%^ cells using NAD(P)H fluorescence (**Extended Data Figure 3D**). Taken together, these data demonstrate that truncating mutations in *mt-Nd5* exert heteroplasmy-dependent effects on the abundance of complex I. In turn, partial loss of complex I disrupts cellular redox balance, without significantly impacting cellular energy homeostasis, oxygen consumption or proliferation.

**Figure 2:**
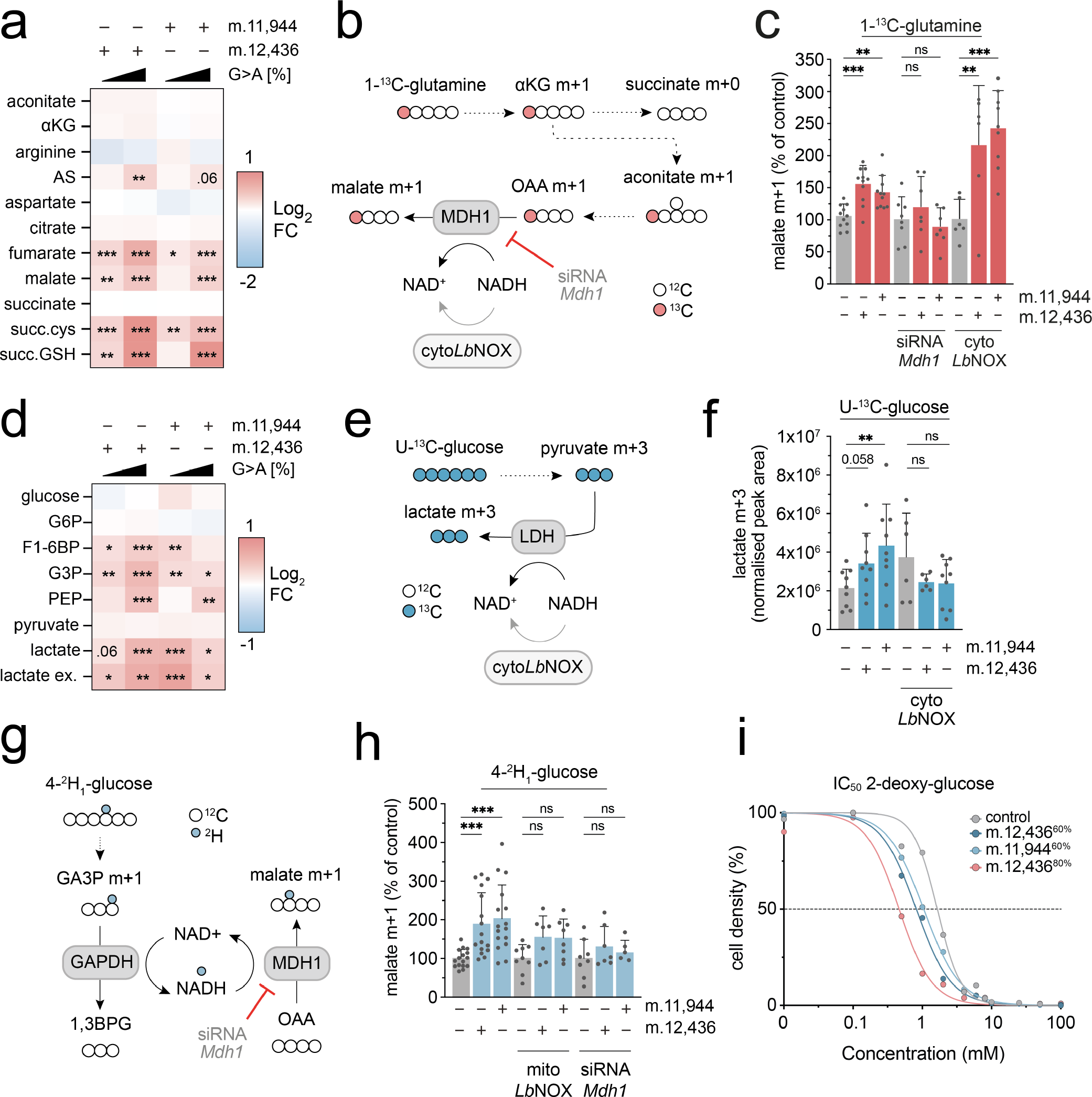
Mutant cells undergo a metabolic shift towards glycolysis due to cellular redox imbalance. **A** Heatmap of unlabelled steady-state abundance of select mitochondrial metabolites, arginine, argininosuccinate (AS) and terminal fumarate adducts succinylcysteine (succ. Cys) and succinicGSH (succ.GSH). **B** Labelling fate of ^13^C derived from 1-^13^C-glutamine. **C** Malate m+1 abundance, derived from 1-^13^C- glutamine with indicated treatment (n = 6-11 separate wells were sampled). **D** Heatmap of unlabelled steady-state metabolite abundances for select intracellular glycolytic intermediates and extracellular lactate (ex. lactate). **E** Labelling fate of U- ^13^C-glucose. **F** Abundance of U-^13^C-glucose derived lactate m+3 with indicated treatment (n = 6-9 separate wells were sampled). **G** Labelling fate of ^2^H derived from 4-^2^H1-glucose; mitoLbNOX not shown for clarity. **H** Malate m+1 abundance, derived from 4-^2^H1-glucose with indicated treatment (n = 5-16 separate wells were sampled). **I** IC50 curves for 2-DG (n = 4 separate wells measured per drug concentration). This was repeated 3 times and a representative result is shown. P-values were determined using a one-way ANOVA test with (A, D) Sidak multiple comparisons test or Fisher’s LSD Test (C, F, H). Error bars indicate SD. Measure of centrality is mean.

**Figure 3:**
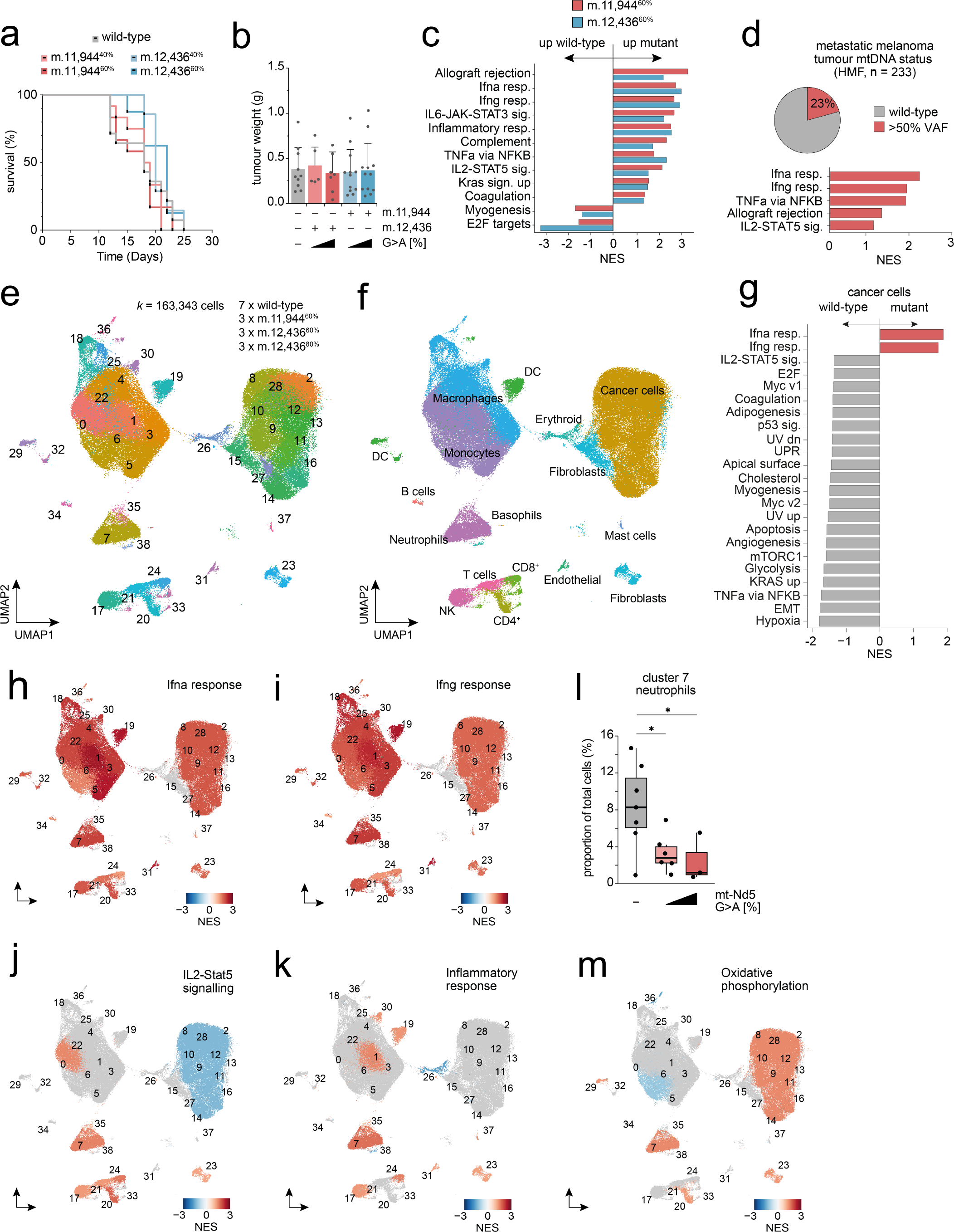
Tumour mtDNA mutations reshape the immune microenvironment. **A** Survival of C57/BL6 mice subcutaneously injected with indicated cells (n = 5-12 animals per condition). **B** Tumour weight at endpoint (n = 5-12 tumours per genotype). **C** Geneset enrichment analysis (GSEA) of bulk tumour RNA sequencing (RNAseq) data (n=5-6 tumours per genotype). Only genesets with adj. P-value <0.1 are shown. **D** GSEA of RNAseq obtained from Hartwig Medical Foundation (HMF) metastatic melanoma patient cohort. Cancers are stratified by mtDNA status into wild-type and mtDNA mutant with >50% variant allele frequency (VAF). **E** UMAP of seurat clustered whole tumour scRNAseq from indicated samples. **F** UMAP indicating cell type IDs. DC, dendritic cells. pDC, plasmacytoid dendritic cell. **G** GSEA of malignant cells identified in scRNAseq analysis. UMAPs coloured by GSEA score for: **H** interferon alpha response; **I** interferon gamma response; **J** inflammatory response; **K** IL2-Stat5 signalling. **L** Proportion of tumour resident neutrophils relative to total malignant and non-malignant cells (n = 17 tumours). **M** UMAP coloured by GSEA for OXPHOS geneset. One-way ANOVA test with Sidak multiple comparisons test (B), Wilcoxon signed rank test (G-K) and two-tailed student’s t-test (L-O) were applied. Error bars indicate SD (B) or SEM (L-O). Measure of centrality is mean. Box plots indicate interquartile range (J-M). NES: normalised expression score.

Unlabelled metabolomic measurements from m.12,436^60%^ and m.11,944^60%^ cells revealed consistent differences in metabolite abundance in these cells relative to control (**Extended Data Figure 4**), with notable increases in the steady-state abundance of malate, lactate, fumarate, argininosuccinate (AS) and the metabolically terminal fumarate adducts succinylcysteine and succinicGSH (**Figure 2A**). Heteroplasmy-dependent increases in abundance of lactate and malate in the context of constant succinate in mutant cells suggested that the flow of electrons into mitochondria through the malate-aspartate shuttle (MAS) might be impacted by changes to the redox state of the cell. To study this we first measured the contributions of glutamine-derived carbon to tricarboxylic acid (TCA) cycle metabolites using U-^13^C- glutamine isotope tracing (**Extended Data Figure 5A)**. This indicated increased abundance of malate from cytosolic oxaloacetate (OAA), derived from citrate via ATP citrate lyase, as determined by the abundance of malate m+3 and the ratio of malate m+3 : m+2, which demonstrated a significant, heteroplasmy-dependent increase relative to control (**Extended Data Figure 5B, C**), with a similar pattern of m+3 : m+2 labelling observed for urea cycle metabolite AS (**Extended Data Figure 5D**). We then traced the metabolic fate of carbon from 1-^13^C-glutamine, which exclusively labels metabolites derived from reductive carboxylation (RC) of glutamine (**Figure 2B, Extended Data Figure 6A).** This revealed that the increased abundance of malate m+1 occurred at the level of MDH1 (**Figure 2C**), but was not apparent in downstream or upstream metabolites aconitate and aspartate (**Extended Data Figure 6B, C**), with the m+1 labelling pattern of AS again matching that of malate (**Extended Data Figure 6D**). The increased abundance of malate m+1 and AS +1 was sensitive to siRNA mediated depletion of *Mdh1* but not expression of cytosolically targeted *Lb*NOX (cyto*Lb*NOX), a water-forming NADH oxidase^10^ (**Figure 2C, Extended Data Figure 6E-G**), indicating that increases in malate abundance occur at least partially in the cytosol via MDH1, but are not directly due to gross alteration in cytosolic NAD+ : NADH redox poise.

**Figure 4:**
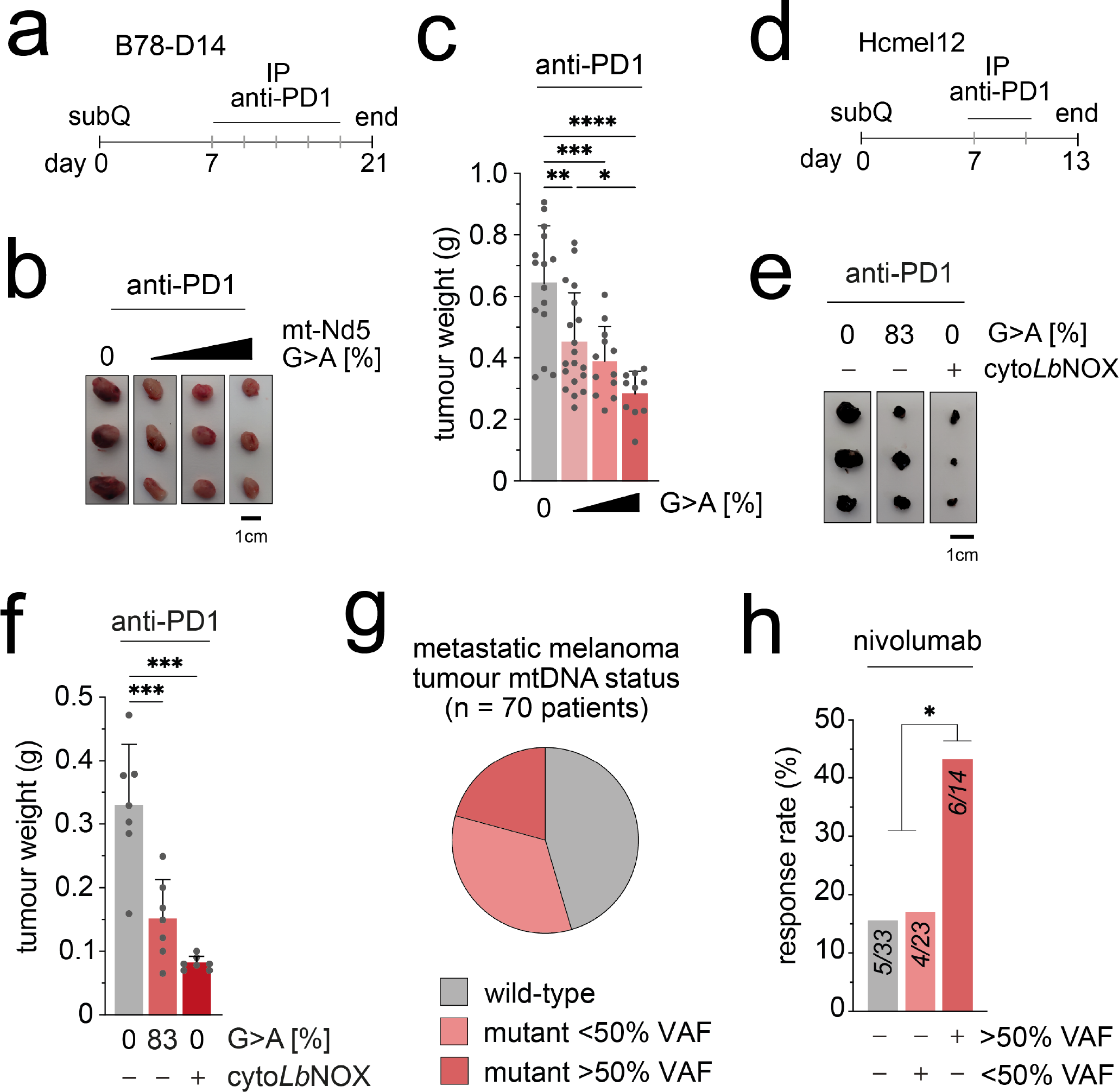
mtDNA mutation-associated microenvironment remodelling sensitises tumours to checkpoint blockade. **A** Schematic of the experimental plan and dosing regimen for B78-D14 tumours with anti-PD1 monoclonal antibody (mAb). **B** Representative images of harvested tumours at day 21. **C** Tumour weights at day 21 (n = 10-19 tumours per genotype). **D** Schematic of experimental plan and dosing regimen for Hcmel12 tumours with anti-PD1 mAb. **E** Representative images of harvested tumours at day 13. **F** Tumour weights at day 13 (n = 7 tumours per genotype). **G** Stratification of a metastatic melanoma patient cohort by mtDNA status. **H** Response rate of patients to nivolumab by tumour mtDNA mutation status. One- way ANOVA test with Sidak multiple comparisons test (C), student’s one-tailed t-test (F)or chi-squared test (H) were applied. Error bars indicate SD. Measure of centrality is mean.

Elevated cellular and extracellular lactate, alongside increased abundance of several glycolytic intermediates (**Figure 2D**) suggested utilisation of pyruvate as an electron acceptor to rebalance NAD+ : NADH via lactate dehydrogenase (LDH). Using U-^13^C-glucose tracing (**Figure 2E**) we observed increased abundance of lactate m+3 in m.12,436^60%^ and m.11,944^60%^ cells that was abolished by cyto*Lb*NOX expression (**Figure 2F, Extended Data Figure 7A**). The increase in lactate m+3 did not alter pyruvate m+3 levels (**Extended Data Figure 7B**), or the entry of glucose-derived carbon into the TCA cycle via pyruvate dehydrogenase (PDH) determined by the ratio of citrate m+2 : pyruvate m+3 (**Extended Data Figure 7C)**. However, the fate of carbon entering the TCA cycle via pyruvate carboxylase (PC) was substantially altered, with a malate m+3 : citrate m+3 ratio indicative of MDH2 reversal (**Extended Data Figure 7D**). Coupling of the MAS with glycolysis is a topic of recent interest, with several reports linking mitochondrial dysfunction with NADH shuttling between GAPDH and MDH1/LDH^11, 12^. Using 4-^2^H1-glucose isotope tracing (**Figure 2G**) we observed an increase in abundance of malate m+1 in m.12,436^60%^ and m.11,944^60%^ cells, with a similar trend in lactate m+1 abundance, that was sensitive to mito*Lb*NOX treatment and siRNA mediated depletion of *Mdh1* (**Figure 2H, Extended Data Figure 8A, B**), supporting the notion that the NAD+ : NADH imbalance resulting from partial loss of complex I supports enhanced glycolytic flux by coupling cytosolic components of the MAS with glycolysis. In turn, this increased glycolytic flux rendered m.12,436^60%^ (IC50= 0.81mM ±0.064mM) and m.11,944^60%^ cells (IC50= 1.04mM ±0.040mM) more sensitive to the competitive phosphoglucoisomerase inhibitor 2-deoxyglucose (2-DG) compared with wild-type cells (IC50= 1.62mM ±0.063mM) (**Figure 2I**), a sensitivity that was further enhanced in a m.12,436^80%^ model (IC50= 0.46mM ±0.080mM). m.12,436^60%^, m.12,436^80%^ and m.11,944^60%^ cells also demonstrated enhanced sensitivity to the low affinity complex I inhibitor metformin relative to wild-type (**Extended Data Figure 9A**). The 60% mutants were not differentially sensitive to potent complex I inhibitor rotenone, although interestingly the m.12,436^80%^ demonstrated resistance compared to wild type (**Extended Data Figure 9B**). None of the mutants demonstrated differential sensitivity to complex V inhibitor, oligomycin (**Extended Data Figure 9C**). Taken together, these data demonstrate that truncating mutations in *mt-Nd5* of complex I induce a Warburg-like metabolic state through redox imbalance, not energetic crisis. This influences both cytosolic and mitochondrial components of the MAS, increasing glycolytic flux, enhancing sensitivity to inhibition of this adaptive metabolic strategy and producing elevated levels of characteristic terminal fumarate adducts succinicGSH and succinylcysteine.

Having established specific changes in redox metabolism driven by truncating mutations in complex I, we next sought to determine the impact of these metabolic alterations in tumour biology. Syngeneic allografts of m.11,944 G>A cells, m.12,436 G>A cells and wild-type controls were performed subcutaneously in immunocompetent C57/Bl6 mice, establishing tumours in 100% of engraftments. All tumours grew at a rate that reached similar humane endpoints (**Figure 3A**) with similar weights and macroscopic histological features (**Figure 3B, Extended Data Figure 10A-C**). Bulk measurements of tumour heteroplasmy revealed a subtle, comparable decrease in heteroplasmy of ∼10% between engrafted cells and resulting tumours, likely reflecting stroma and immune cell infiltrate (**Extended Data Figure 10D**), with no consistent change in mtDNA copy number detected at bulk level (**Extended Data Figure 10E**). Measurements of metabolites from m.11,944^60%^ mutant and control tumours revealed elevated abundance of terminal fumarate adducts succinicGSH and succinylcysteine, characteristic of the metabolic rewiring observed *in vitro* (**Extended Data Figure 10F**). These markers of a consistently altered tumour metabolic profile were coupled to divergent transcriptional signatures between control and mutant tumours (**Figure 3C**), with several signatures of altered immune infiltrate and signalling being significantly elevated in mutant tumours compared with controls, notably allograft rejection, interferon gamma (Ifng) and interferon alpha (Ifna) responses. Higher heteroplasmies correlated to increased signal in the same gene sets (**Extended Data Figure 11**) suggesting a heteroplasmy dose-dependent anti-tumour immune response. To benchmark these findings against human data, we took the Hartwig Medical Foundation (HMF) metastatic melanoma cohort and stratified this by pathogenic mtDNA mutation status into wild-type and >50% variant allele frequency (VAF) groups (see **Methods**). This yielded a set of 355 tumour samples (272 wildtype, 83 >50% VAF), with 233 having transcriptional profiles. GSEA analysis revealed consistent transcriptional phenotypes between patient tumours bearing high heteroplasmy pathogenic mtDNA mutations and those identified in our model systems (**Figure 3D**), supporting the observation. To further dissect these effects we employed whole tumour single cell RNA sequencing (scRNAseq) across seven control, three m.12,436^60%^, three m.11,944^60%^ and three m.12,436^80%^ tumours, resulting in 163,343 single cell transcriptomes. Cells were clustered using Seurat and cellRanger, with preliminary cell ID determined by scType (see **Methods**) (**Figure 3E,F**). Malignant cells were assigned on the basis of: i) low or nil *Ptprc* (CD45) expression; ii) high epithelial score^13^; iii) aneuploidy determined by copykat analysis^14^ (**Extended Data Figure 12**). Consistent with bulk tumour transcriptional profiles, GSEA in malignant cells revealed increased Ifna and Ifng signatures coupled to decreased glycolysis signatures in high heteroplasmy tumours (**Figure 3G**), which is not observed *in vitro* prior to implantation (**Extended Data Figure 13**). Downstream regulation of primary metabolic and subsequent immune signalling on malignant cells are also reflected in altered nutrient sensing by mTORC1, transcriptional control of metabolic genes by myc, and TNFa signalling (**Figure 3G)**. GSEA in non-malignant cell clusters revealed similar tumour-wide changes in transcriptional phenotype, with increased Ifna, Ifng, inflammatory response and IL2-Stat5 signalling again observed (**Figure 3H-K**). These indicators of a broad anti-tumour immune response were accompanied by decreased neutrophil residency (**Figure 3L**) and altered monocyte maturation (**Extended Data Figure 14A, B)**, with a switch in neutrophil metabolic state indicated by increased OXPHOS gene expression (**Figure 3M**). Further genesets typical of an augmented anti-tumour response, such as allograft rejection, were also elevated alongside a biphasic trend in proportions of tumour resident natural killer and CD4+ T cells (**Extended Data Figure 14C-E**). Taken together these data demonstrate that, in a heteroplasmy-dependent fashion, *mt-Nd5* mutation is sufficient to remodel the tumour microenvironment (TME) and promote an anti-tumour immune response.

Treatment of malignant melanoma can include immune checkpoint blockade (ICB) with monoclonal antibodies (mAbs) against T and B-cell expressed immune checkpoint receptor PD1, blocking PD-L1/2 binding to limit tumour-induced immune tolerance. However, the effectiveness of anti-PD1 treatments, and ICB response in melanoma patients more broadly is bimodal, with a substantial proportion of patients not responding to treatment while experiencing a poor morbidity profile. Limited efficacy of ICB has been linked to immunosuppressive tumour-associated neutrophils previously^15^ therefore we reasoned that *mt-Nd5* mutant tumours could demonstrate differential sensitivity to ICB, even in an aggressive model of poorly immunogenic melanoma such as B78-D14. Additionally, depleted neutrophil populations in *mt-Nd5* mutant tumours also demonstrated the greatest PD-L1 expression (**Extended Data Figure 14F**). To test this we performed further subcutaneous syngeneic allografts of m.12,436^40%^, m.12,436^60%^, m.12,436^80%^, m.11,944^40%^, m.11,944^60%^ and wild-type tumours in immunocompetent animals. Tumours grew untreated for 7 days post-graft and animals were dosed with a regimen of intraperitoneal anti-PD1 mAb every 3 days until conclusion of the experiment (**Figure 4A**). A heteroplasmy-defined decrease in tumour weight at endpoint was observed across the mtDNA mutant tumours, with higher mutant heteroplasmies exhibiting greater response to treatment (**Figure 4B,C, Extended Figure 15**), consistent with increased sensitivity of mtDNA mutant tumours to immunotherapy. To verify these data we attempted to produce further independent models of aggressive, poorly immunogenic mouse melanoma (**Extended Data Figure 16A**). This yielded Hcmel12 (Hgf, Cdk4^R24C^)^16^ cells engineered to bear >80% m.12,436 G>A mutation, demonstrating consistent cellular and metabolic phenotypes with B78-D14 (**Extended Data Figure 16B-J)**. Hcmel12 m.12,436^80%^ and wild-type Hcmel12 cells were engrafted into mice with a similar experimental workflow as previously (**Figure 4D**). When untreated, Hcmel12 m.12,436^80^ and wild-type tumours demonstrate comparable time to endpoint and tumour weight at endpoint (**Extended Data Figure 17A, B**). Changes in bulk heteroplasmy, copy number and tumour metabolism were also similar to those of B78-D14 tumours (**Extended Data Figure 17C-D).** Moreover, when anti-PD1 treatment was administered, a mtDNA mutation-dependent response was observed in Hcmel12 of similar magnitude to that seen in B78-D14 (**Figure 4E,F**). To dissect the enhanced ICB response into metabolic v.s. non-metabolic effects of mtDNA mutation, we modified wild-type Hcmel12 cells to constitutively express cyto*Lb*NOX, which reproduces key elements of the cell-extrinsic, mutant *Mt-Nd5*-associated metabolic phenotype, notably glucose uptake and lactate release (**Extended Data Figure 18**). When grafted into mice, Hcmel12 cyto*Lb*NOX tumours demonstrated comparable time to endpoint and tumour weight at endpoint as wild-type or *Mt-Nd5* mutant tumours (**Extended Data Figure 17A,B**). When challenged with anti-PD1 treatment, Hcmel cyto*Lb*NOX tumours recapitulate the response of Hcmel *mt-Nd5* m.12,436^80%^ tumours, indicating that specific changes in redox metabolism associated with mtDNA mutation are sufficient to sensitise the tumour to ICB (**Figure 4E,F**). To benchmark these findings from mice against real world clinical data, we re-analysed a previously reported, well-characterised cohort of majority treatment-naive metastatic melanoma patients given a dosing regimen of the anti-PD1 mAb nivolumab^17^. By identifying mtDNA mutant cancers and stratifying this patient cohort solely on the basis of cancer mtDNA mutation status (**Figure 4G**) the 70 patients in this cohort were divided into three groups: mtDNA wild-type (33), <50% VAF (23), and >50% VAF (14). The cancer mtDNA mutation status-naive cohort response rate was 22% for partial or complete responses to nivolumab, however the rate of response for >50% mtDNA mutation VAF cancers was 2.6-fold greater than wild-type or <50% VAF cancer (**Figure 4H**), recapitulating our laboratory findings in patients.

These data confirm that somatic mtDNA mutations, commonly observed in human tumours, can exert direct effects on cancer cell metabolic phenotypes. In contrast with clinically presented germline mtDNA mutations,^6^ tumour mtDNA mutations are able to exert these effects at a comparably low heteroplasmic burden and without necessarily negatively impacting oxygen consumption or energy homeostasis. The direct link observed between redox perturbations and enhanced glycolytic flux subtly alters our view of mtDNA mutation, to an adaptive gain of function rather than exclusively loss of function event, and the discovery that mtDNA mutations can underpin aerobic glycolysis warrants further assessment of the relationship between classical Warburg metabolism^18^ and mtDNA mutation status.

Beyond cancer cell intrinsic effects, the data here reveal that a functional consequence of somatic mtDNA mutation in tumour biology is the remodelling of the TME, mediating therapeutic susceptibility to ICB. Truncating mutations to mtDNA, analogous to those described here, affect 10% of all cancers regardless of tissue lineage, with non-truncating, pathogenic mtDNA mutations presenting in a further 40- 50% of all cancers. A broad influence over the anti-tumour immune response in these cancers might also be expected.

Beyond stratification and exploitation of mtDNA mutant tumour vulnerability, our data suggest that the ICB response-governing effects we observe are principally metabolic in nature. Recreating such a metabolic state in mtDNA wild-type or ‘immune cold’ tumour types could therefore also be of benefit.

## Methods

### Maintenance, transfection and FACS of cell lines

B78 melanoma cells (RRID:CVCL_8341) and Hcmel12 cells^16^ were maintained in DMEM containing GLUTAMAX^TM^, 0.11g/L sodium pyruvate, 4.5g/L D-glucose (Life Technologies) and supplemented with 1% penicillin/ streptomycin (P/S) (Life Technologies) and 10% FBS (Life Technologies). Cells were grown in incubators at 37°C and 5% CO2. Cells were transfected using Lipofectamine 3000 (Life Technologies) using a ratio of 5µg DNA : 7.5µl Lipofectamine 3000. Cells were sorted as outlined in^19^ and thereafter grown in the same base DMEM media supplemented with 20% FBS and 100µg/mL of uridine (Sigma).

### Use of animal models

Animal experiments were carried out in accordance with the UK Animals (Scientific Procedures) Act 1986 (P72BA642F) and by adhering to the ARRIVE guidelines with approval from the local Animal Welfare and Ethical Review Board of the University of Glasgow. Mice were housed in conventional cages in an animal room at a controlled temperature (19–23 °C) and humidity (55 ± 10%) under a 12hr light/dark cycle. Experiments only used male C57BL/6 mice at ∼8 weeks of age which were injected subcutaneously with either 2.5x10^5^ B78 cells or 1x10^4^ HcMel12 cells, both prepared in 1:1 RPMI (Life Technologies) and Matrigel (Merck). Mice were culled at an endpoint of 15mm tumour measurement.

For immunotherapy experiments, mice were put on a dosing regimen of 200µg of anti- PD1 given intraperitoneally twice a week. The first dose was given 7 days post-injection and all mice were sacrificed at 21 or 13 days post-injection for B78 or HcMel12 cells respectively.

### Construction of DdCBE plasmids

TALEs targeting mt.12,436 and mt.11,944 were designed with advice from Beverly Mok and David Liu (Broad Institute, USA). TALEs were synthesised (ThermoFisher GeneArt) as illustrated in Figure 1A with the left TALEs being cloned into pcDNA3.1(-)_mCherry^19^ and the right into pTracer CMV/Bsd^19^, allowing for the co-expression of mCherry and GFP respectively.

### Pyrosequencing assay

DNA was extracted from cell pellets using the DNeasy Blood & Tissue Kit (Qiagen) as per the manufacturer’s instructions. PCR was then performed using the PyroMark PCR Mix (Qiagen) for 50 cycles with an annealing temperature of 50°C and an extension time of 30sec. PCR products were run on the PyroMark Q48 Autoprep (Qiagen) as per the manufacturer’s instructions.

PCR primers for mt.12,436

Forward: 5’-ATATTCTCCAACAACAACG-3’ Reverse: 5’-**biotin**-GTTATTATTAGTCGTGAGG-3’

PCR primers for mt.11,944

Forward: 5’-CTTCATTATTAGCCTCTTAC-3’ Reverse: 5’-**biotin**-GTCTGAGTGTATATATCATG-3’

Sequencing primer for mt.12,436 5’-TTGGCCTCCACCCAT-3’

Sequencing primer for mt.11,944 5’-TAATTACAACCTGGCACT-3’

### Protein extraction and measurement

Cell pellets were lysed in RIPA buffer (Life Technologies) supplemented with cOmplete Mini Tablets and cOmplete Mini Protease Inhibitor Tablets (Roche). Samples were incubated on ice for 20mins and then spun at 14,000*g* for 20mins. The isolated supernatant containing total cellular protein was then quantified using a DC Protein Assay (Bio-Rad Laboratories) performed as per the manufacturer’s instructions.

### Immunoblotting

To detect protein via western blotting 60µg of protein was resolved on SDS-PAGE 4- 12% Bis-Tris Bolt gels (Life Technologies). Protein was transferred onto a nitrocellulose membrane using a Mini Trans-Bolt Cell (Bio-Rad Laboratories). Membranes were then stained with Ponceau S Staining Solution (Life Technologies) to measure loading before overnight incubation with the primary antibody prepared in 5% milk in 1X TBST. Imaging was performed using the Odyssey DLx Imaging system (Licor).

Antibodies:

Total OXPHOS Rodent WB Antibody Cocktail (1:800, ab110413, Abcam) Monoclonal Anti-FLAG^®^ M2 antibody (1:1000, F1804, Sigma) Recombinant anti-vinculin antibody (1:10,000, ab129002, Abcam)

### Mitochondrial Isolation

Cells were grown in Falcon Cell Culture 5-layer flasks (Scientific Laboratory Supplies) and grown to near 100% confluency. Cells were then harvested and mitochondria were extracted as described in ^20^.

### Blue-Native PAGE

Isolated mitochondria were solubilized in 1X NativePage Sample Buffer supplemented with 1% Digitonin (Life Technologies). Samples were incubated on ice for 10min and then centrifuged at 20,000*g* for 30min at 4°C. Supernatants were isolated and total extracted protein quantified using the DC Protein Assay (Bio-Rad Laboratories). Samples were prepared and run on NativePage 4-12% Bis-Tris gels as per the manufacturer’s instructions (Life Technologies). For immunoblotting, samples were transferred onto PVDF membranes using Mini Trans-Bolt Cell (Bio-Rad Laboratories). Subsequent probing and imaging was performed as described above for immunoblotting. Loading was visualised using Coomassie Blue on a duplicate gel.

In-gel assays were performed for complex I and II activity as described in ^20^.

### Digital droplet PCR mt-Nd5 primers

Forward: 5’-TGCCTAGTAATCGGAAGCCTCGC-3’ Reverse: 5’-TCAGGCGTTGGTGTTGCAGG-3’

VDAC1 primers

Forward: 5’-CTCCCACATACGCCGATCTT-3’

Reverse: 5’-GCCGTAGCCCTTGGTGAAG-3’

Samples were prepared in triplicate in a 96-well plate using 1ng of DNA, 100nM of each primer, 10µL of QX200 ddPCR EvaGreen Supermix and water to 20µL. Droplet generation, PCR and measurements were then performed on the QX200 Droplet Digital PCR System (Bio-Rad Laboratories) as per the manufacturer’s instructions with the primer annealing temperature set at 60°C.

### Seahorse Assay

The Seahorse XF Cell Mito Stress Test (Agilent) was performed as per the manufacturer’s instructions. Briefly, cells were plated into a Seahorse 96-well plate at 2 x 10^4^ cells/well a day prior to the assay. A sensor cartridge was also allowed to hydrate in water at 37°C overnight. The water was replaced with Seahorse XF Calibrant and the sensor cartridge was re-incubated for 45mins. Oligomycin, FCCP, Rotenone and Antimycin A were then added to their respective seahorse ports to a final concentration of 1µM in the well before sensor calibration on the Seahorse XFe96 Analyser (Agilent). Meanwhile, cell media was replaced with 150µL Seahorse XF Media supplemented with 1% FBS, 25mM glucose, 1mM sodium pyruvate and 2mM glutamine and incubated at 37°C for 30mins. The cell plate was then inserted into the analyser post-calibration and run.

For read normalisation, protein extraction and measurement was performed as described above.

### In vitro metabolomics

Cells were seeded two days prior to metabolite extraction to achieve 70-80% confluency on the day of extraction. Plates were incubated at 37°C and 5% CO2 overnight. The following day, cells were replenished with excess fresh media to prevent starvation at the point of extraction. For steady-state experiments, media was prepared as described above with the substitution of GLUTAMAX™ with 2mM L-glutamine. For U-^13^C-glucose and 4-^2^H1-glucose isotope tracing experiments, media was prepared as follows: DMEM, no glucose (Life Technologies) supplemented with 0.11g/L sodium pyruvate, 2mM L-glutamine, 20% FBS, 100µg/mL uridine and 25mM glucose isotope (Cambridge Isotopes). For isotope tracing experiments using U-^13^C- glutamine and 1-^13^C-glutamine, DMEM containing 4.5g/L D-glucose and 0.11g/L sodium pyruvate was supplemented with 20% FBS, 100µg/mL uridine and 4mM glutamine isotope (Cambridge Isotopes).

On the day of extraction, 20µL of media was added to 980µL of extraction buffer from each well. Cells were then washed twice with ice-cold PBS. Extraction buffer (50:30:20, v/v/v, methanol/acetonitrile/water) was then added to each well (600µL per 2 x10^6^) and incubated for 5min at 4°C. Samples were centrifuged at 16,000*g* for 10mins at 4°C and the supernatant was transferred to liquid chromatography-mass spectrometry (LC-MS) glass vials and stored at -80°C until run on the mass spectrometer.

Mass spectrometry and subsequent targeted metabolomics analysis was performed as described in ^21^. Compound peak areas were normalised using the total measured protein per well quantified with a modified Lzowry assay^21^.

### In vitro measurements of fumarate

#### Samples were prepared as described above

Fumarate analysis was carried out using a Q Exactive Orbitrap mass spectrometer (Thermo Scientific) coupled to an Ultimate 3000 HPLC system (Themo Fisher Scientific). Metabolite separation was done using a HILIC-Z column (InfinityLab Poroshell 120, 150 x 2.1 mm, 2.7µm, Agilent) with a mobile phase consisting of a mixture of A (40mM ammonium formate pH=3) and B (90% ACN / 10% 40 mM ammonium formate). The flow rate was set to 200 µL/min and the injection volume was 5 µL. The gradient started at 10% A for 2 min, followed by a linear increase to 90% A for 15 min; 90% A was then kept for 2 minutes, followed by a linear decrease to 10% A for 2 min and a final re-equilibration step with 10% A for 5 min. The total run time was 25 min. The Q Exactive mass spectrometer was operated in negative mode with a resolution of 70,000 at 200 *m/z* across a range of 100 to 150 *m/z* (automatic gain control (AGC) target of 1x10^6^ and maximum injection time (IT) of 250 ms).

### siRNA knockdown for metabolomics

1.2 x 12^4^ cells were plated into 12-well cell culture plates and incubated at 37°C and 5% CO2 overnight. The following day, cells were transfected with 5µL of 5µM siRNA with 5µL of DharmaFECT 1 Transfection Reagent (Horizon Discovery). Cells were either transfected with ON-TARGETplus MDH1 siRNA (L-051206-01-0005, Horizon Discovery) or ON-TARGETplus non-targeting control siRNA (D-001810-10-05, Horizon Discovery). Cells were supplemented with excess media the following day and metabolites extracted 48hrs post-transfection as outlined above.

### LbNOX treatment for metabolomics

pUC57-LbNOX (addgene #75285) and pUC57-mitoLbNOX (addgene #74448) were gifts from Vamsi Mootha. Both enzyme sequences were amplified using Phusion PCR (Life Technologies) as per the manufacturer’s instructions. These products were cloned into pcDNA3.1(-)_mCherry^19^ via the *NheI* and *BamHI* restriction sites and used for subsequent experiments.

Forward for LbNOX: 5’-GGTGGTGCTAGCCGCATGAAGGTCACCG-3’ Forward for mitoLbNOX: 5’-GGTGGTGCTAGCCGCATGCTCGCTACAAG-3’ Reverse: 5’-GGTGGTGGATCCTTACTTGTCATCGTCATC-3’

Cells were transfected and sorted as described above and 3 x 10^4^ mCherry+ cells were plated per well into a 12-well plate. Cells were allowed to recover overnight at 37°C and 5% CO2 followed by the addition of excess media to each well. Metabolites were extracted the following day and analysed as outlined above.

### Bulk tumour metabolomics

Tumour fragments (20-40mg) were flash frozen on dry ice when harvested. Metabolites were extracted using the Precellys Evolution homogenizer (Bertin) with 25µL of extraction buffer per mg of tissue. Samples were then centrifuged at 16,000g for 10mins at 4°C and the supernatant was transferred to LC-MS glass vials and stored at -80°C until analysis.

Samples were run and subsequent targeted metabolomics analysis was performed as described in ^21^. Compound peak areas were normalised using the mass of the tissue.

### Calculating cell sensitivity to 2-DG

Cells were plated in a 96-well plate at 500 cells/well in 200µL of cell culture media. Plates were incubated overnight at 37°C and 5% CO2. The following day, media was replaced with 0 – 100mM 2-DG in quadruplicate. Plates were imaged once every 4 hrs on the IncuCyte Zoom (Essen Bioscience) for 5 days. Final confluency measurements were calculated using the system algorithm and the IC50 was determined by GraphPad Prism.

### Bulk tumour RNAseq

Tumour fragments (20-40mg) were stored in RNAlater (Sigma) and stored at -80°C. Samples were sent to GeneWiz Technologies for RNA extraction and sequencing.

### HcMel12 Transduction

cyto*Lb*NOX was cloned into the lentiviral plasmid pLex303 via the *NheI* and *BamHI* restriction sites and transduction of HcMel12 was performed as described in ^22^. Transduced cells were selected via supplementation of 8µg/mL blasticidin, and single clones were selected out from the surviving bulk population. cytoLbNOX expression was confirmed using immunoblotting.

pLEX303 was a gift from David Bryant (Addgene plasmid #162032; http://n2t.net/addgene:162032 ; RRID:Addgene_162032).

### Hartwig Dataset Analysis

The Hartwig Medical Foundation (HMF) dataset included WGS data from tumor metastases normal-matched samples from 355 melanoma patients (skin primary tumor location), of whom 233 had additional RNA sequencing data of the tumor samples. mtDNA somatic mutations were called and annotated as previously described^1^.In brief, variants called by both Mutect2 and samtools mpileup were retained and merged using vcf2maf, which embeds the Variant Effect Predictor (VEP) variant annotator. Variants within the repeat regions (chrM:302-315, chrM:513-525, and chrM:3105-3109) were filtered out. Next, variants were filtered out if the Variant Allele Fraction (VAF) was lower than 1% in the tumor samples and lower than 0.24% in the normal sample, as previously described (Yuan et al, 2020). Finally, somatic variants were kept when supported by at least one read in both the forward and the reverse orientations. Samples with >50% VAF mtDNA Complex I truncating (frameshift indels, translation start site and nonsense mutations) and missense mutations were classified as mutated and the rest as wild-type. Gene expression data was obtained from the output generated by the isofox pipeline, provided by HMF. Adjusted Transcript per Million (“adjTPM’’) gene counts per sample were merged into a matrix. Gene expression and mutation data were used to perform differential expression analysis with DESeq2 in R using the DESeqDataSetFromMatrix function. Gene set enrichment analysis (GSEA) was performed with fGSEA in R with a minimum set size of 15 genes, a maximum of 500 genes and 20,000 permutations, against the mSigDB Hallmark gene set collection (v.7.5.1). Normalized Enrichment Score (NES) were ranked for significant upregulated and downregulated gene sets.

#### Statistical methods

No statistical test was used to determine sample sizes. Mice were randomly assigned to different experimental groups. Samples were blinded to machine operators (metabolomics, proteomics, RNAseq). Researchers were blinded to experimental groups for *in vivo* anti-PD1 experiments. Specific statistical tests used to determine significance, group sizes (*n*) and *P* values are provided in the figure legends. *P* values < 0.05, <0.01 and <0.001 are represented as *, ** and *** respectively in figures. All statistical analysis was carried out using Prism (GraphPad) and Rstudio.

#### Data and Code Availability Statement

All non-commercial plasmids used have been deposited with addgene (Gammage Lab). All metabolomic data, mtDNA sequencing, bulk and single cell RNAseq and proteomic data contained in this study are available in the supplementary information or via specified public repositories.

All custom code will be made available via Reznik lab Github.

## Acknowledgements

P.A.G. is grateful to C.Frezza (CECAD, Cologne) E.Chouchani (DFCI, Harvard), O. Sansom (CRUK BI) S. Coffelt (CRUK BI) and J. Norman (CRUK BI) for helpful discussion. The authors would like to acknowledge the advice of B. Mok and D. Liu (Broad Institute) regarding TALE design. This publication and the underlying study have been made possible partly based on data that Hartwig Medical Foundation and the Center of Personalized Cancer Treatment (CPCT) have made available to the study through the Hartwig Medical Database.

## Author contributions

M.M., E.R. and P.A.G. conceived the study. M.M. and P.A.G. designed the experiments. M.M. conducted in vitro and in vivo experiments, analysed data and co-wrote the paper. E.M.L., M.K., T.P. and J.L.M. and E.R. conducted computational analyses. A.S., E.T., A.L.Y. and E.W.R conducted in vivo experiments. J.T.-M. conducted in vitro experiments. A.U., E.S. and D.S. performed metabolomic mass spectrometry. S.L. and S.Z. performed proteomic mass spectrometry and analysis. R.W., R.J.S. and J.N.B performed biophysical experiments. C.R.-A supervised computational analyses. E.R. and P.A.G. supervised the study, obtained funding (CRUK BI Core Funding: A_BICR_1920_Gammage to P.A.G.; ERC Starting Grant via UKRI: EP/X035581/1 to P.A.G.; NIH NCI: 1R37CA276200 to E.R. and P.A.G.) and wrote the paper, with the involvement of all authors.

## Materials & Correspondence

All requests for biological materials, computer code or data should be addressed to

## Extended Data Figures

**Extended Data Figure 1.**
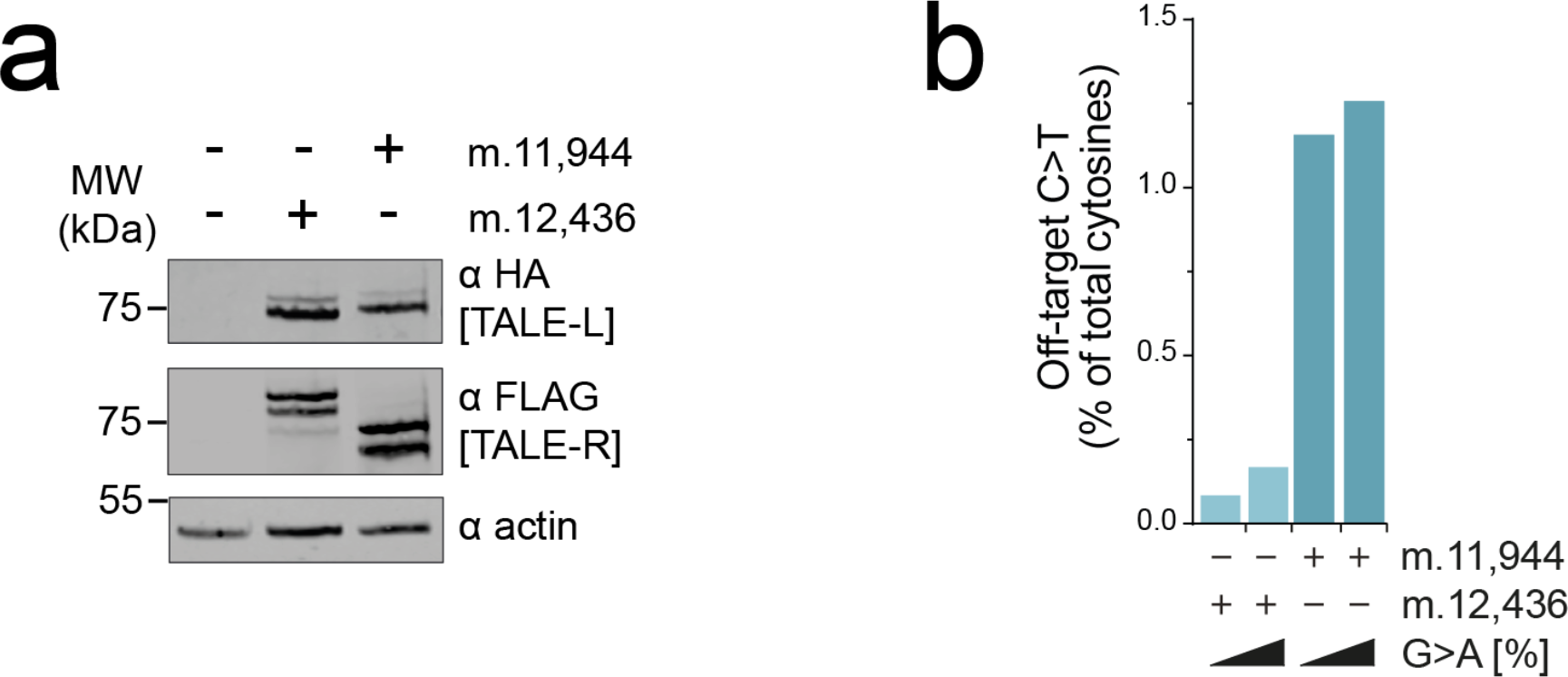
**Mitochondrial base editors for two independent targets in *mt-Nd5***. **A** Immunoblot of DdCBE pair expression post-sort. αHA and αFLAG show expression of left (TALE-L) and right TALEs (TALE-R) respectively. Representative result is shown. **B** Off-target C>T activity of DdCBEs on mtDNA by ultra-deep amplicon resequencing of whole mtDNA. Figure depicts mutations detected at heteroplasmies >2% and is a measure of mutations detected relative to wild-type. These mutations likely do not impact our key observations as both models behave similarly across experiments.

**Extended Data Figure Figure 2.**
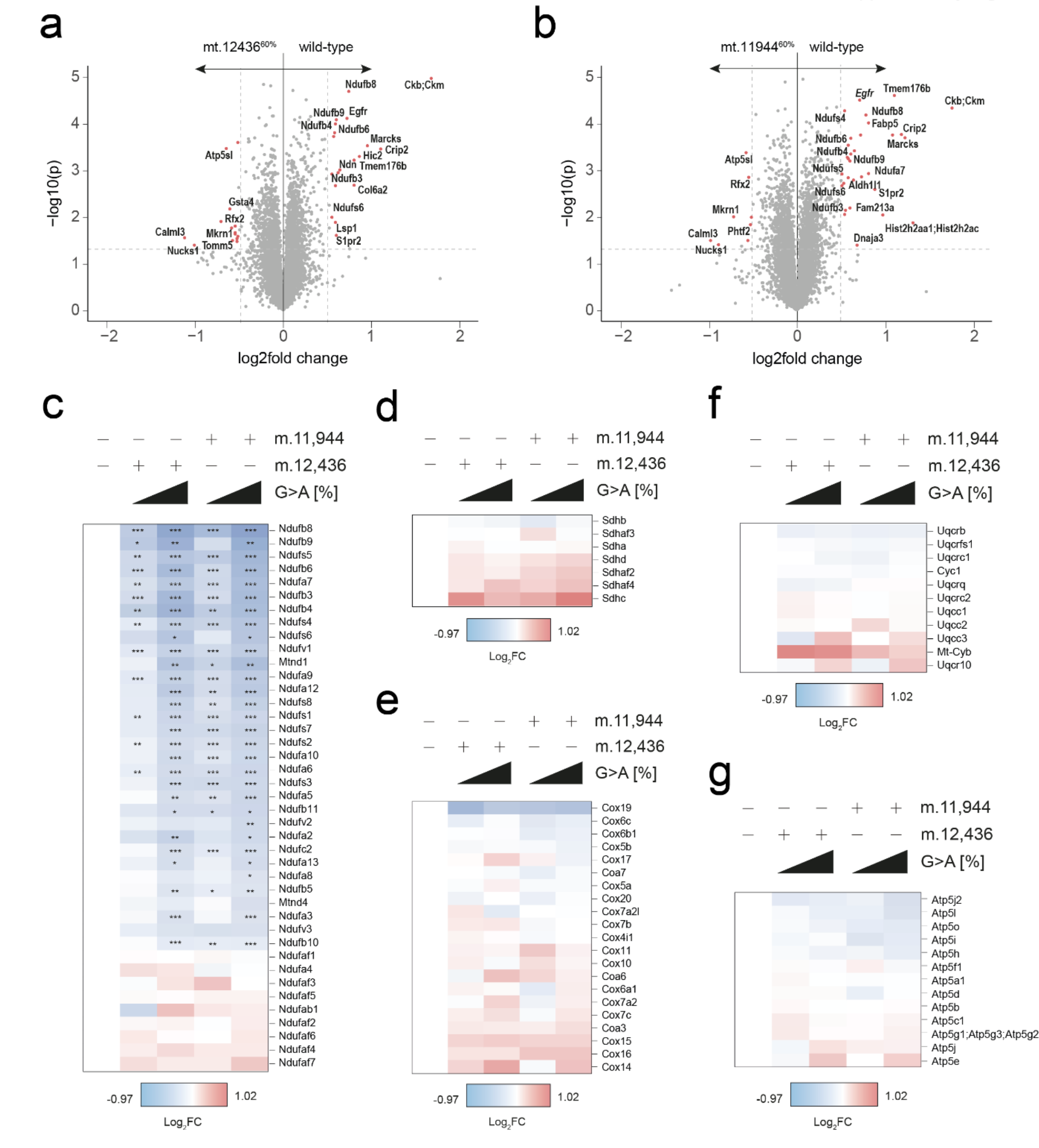
**Proteomic analysis of isogenic *mt-Nd5* mutant cell lines reveals significant changes primarily in complex I genes**. Volcano plot showing detected differences in protein abundance of **A** mt.12436^60%^ cells and **B** mt.11944^60%^ cells versus wild-type. Differences of p < 0.05 and log2 fold change > 0.5 shown in red (n=3 separately collected cell pellets were measured per cell line). Heatmaps of protein abundances for **C** complex I, **D** complex II, **E** complex III, **F** complex IV and **G** complex V nuclear and mitochondrial subunits. Wilcoxon signed rank test (A, B) and a one-way ANOVA test with Sidak multiple comparisons test (C-G) were applied

**Extended Data Figure Figure 3.**
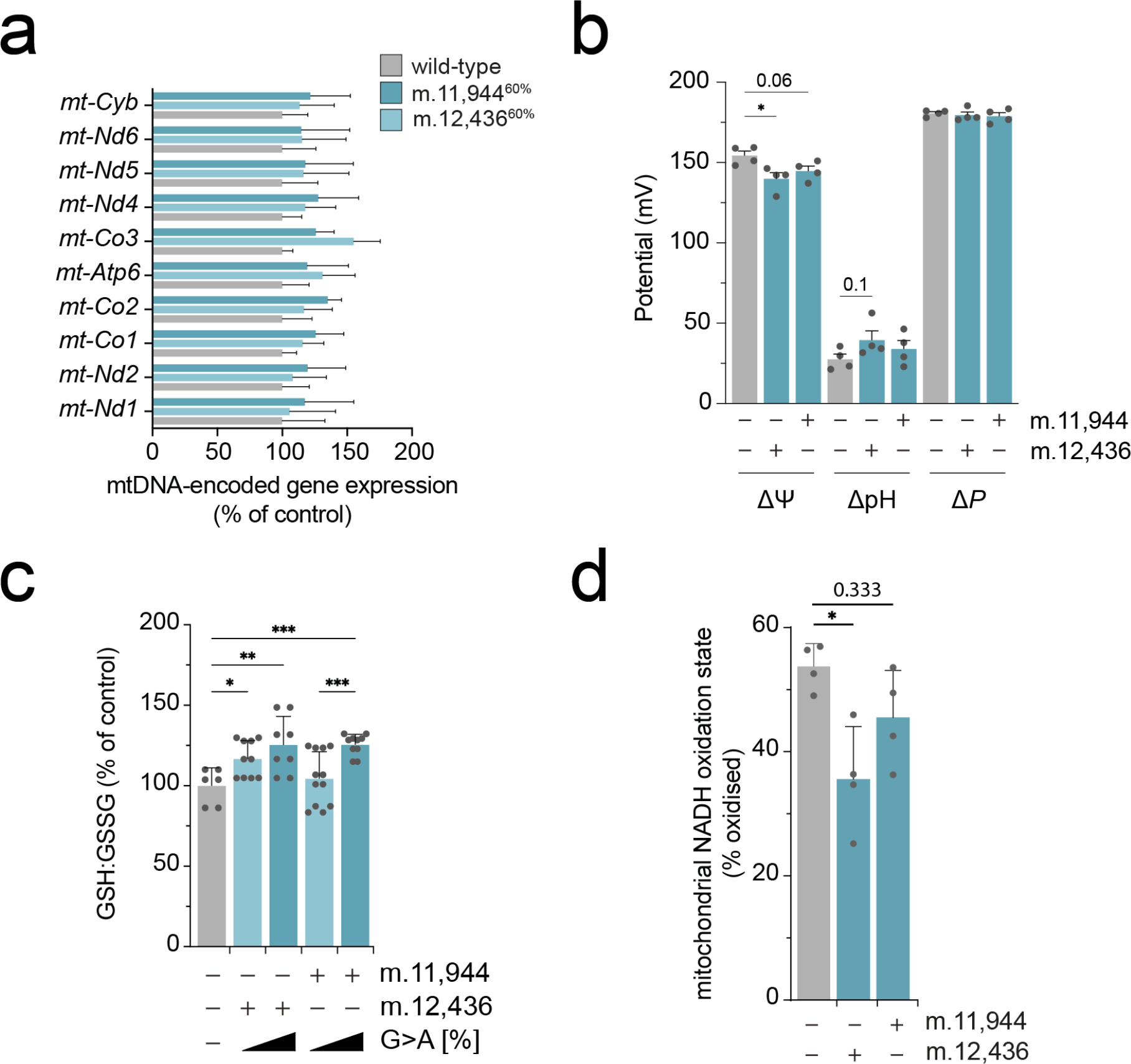
***mt.-Nd5* truncations do not impact mitochondrial mRNA expression levels, but alter intracellular redox state and mitochondrial membrane potential**. **A** Expression of mitochondrial genes (n=12 separate cell pellets were sampled per genotype). **B** Measurements of the electrical component of the proton motive force, Δ^Ψ^, the chemical component of the proton motive force ΔpH and total protonmotive force, Δ*P* (n=4 separate wells were sampled per genotype). **C** GSH : GSSG ratio (n= 6-12 separate wells were sampled per cell type). A high GSH : GSSG ratio represents a more reductive intracellular environment. **D** Mitochondrial NADH oxidation state (n=4 separate wells for sampled per genotype). All P-values were determined using a one-way ANOVA test with Sidak multiple comparisons test. Error bars indicate SD. Measure of centrality is mean.

**Extended Data Figure Figure 4.**
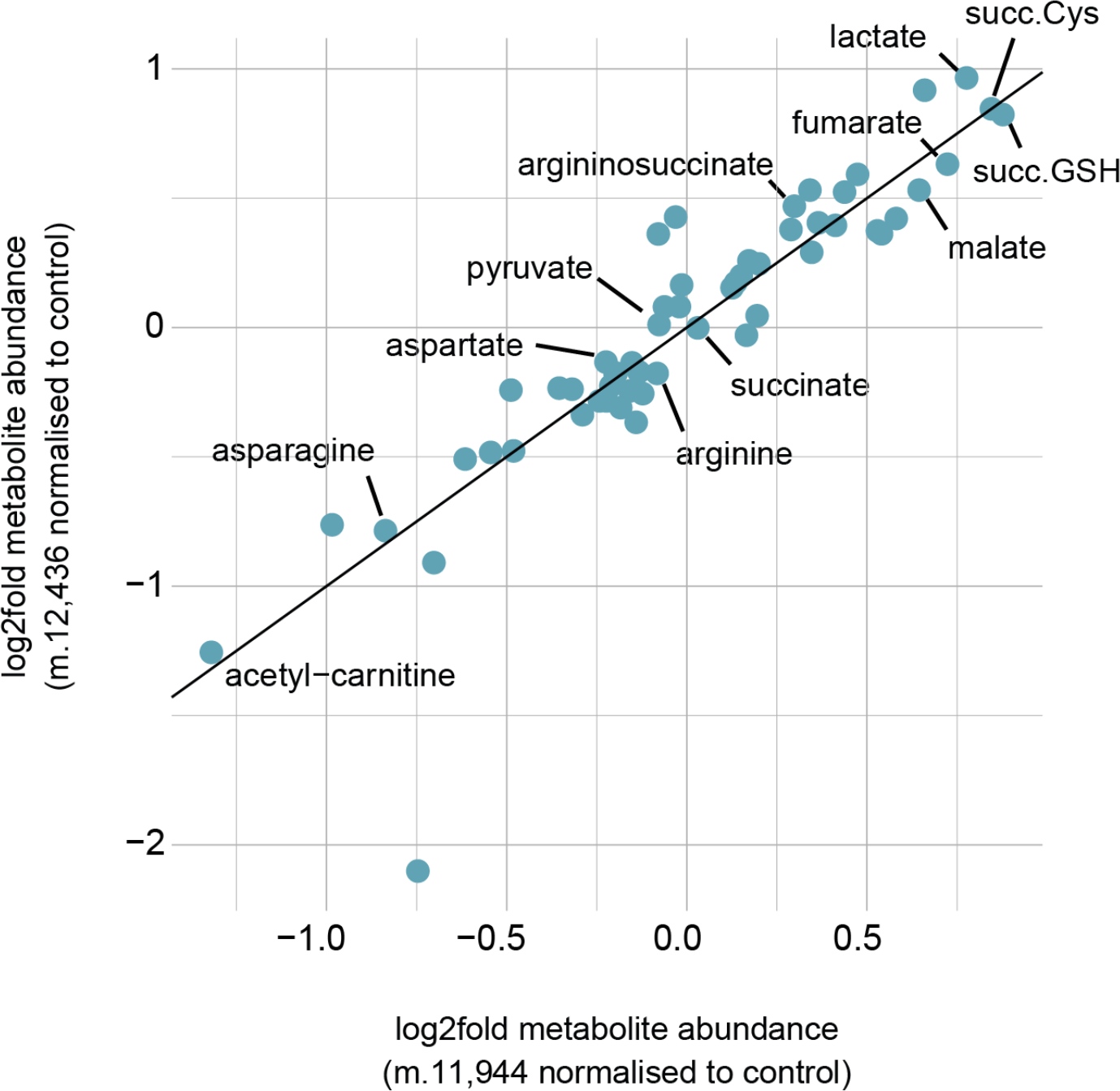
**Independent *mt-Nd5* truncations at matched heteroplasmy produce consonant changes in metabolite abundance**. Comparison of steady-state metabolite changes of m.12,436^60%^ and m.11,944^60%^ cells, each relative to wild-type (n= 6-9 separate wells per sample).

**Extended Data Figure Figure 5.**
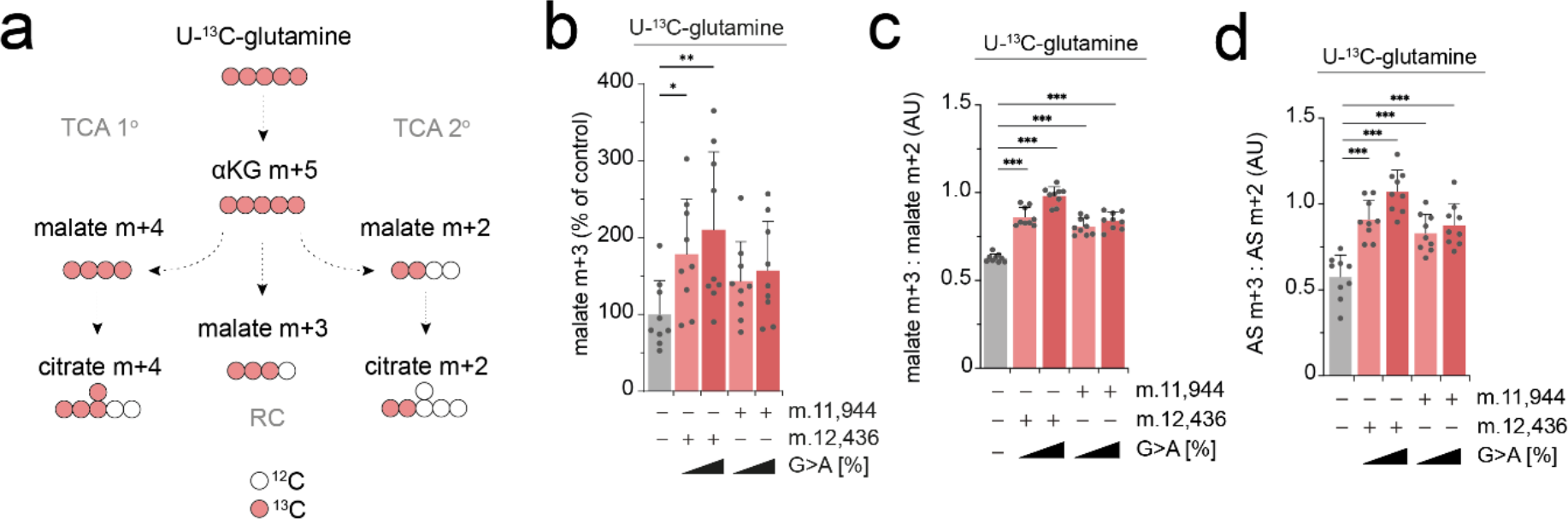
**U-^13^C-glutamine labelling demonstrates that a proportion of the increased malate abundance is derived from cytosolic oxaloacetate**. **A** Labeling fate of ^13^C derived from U-^13^C-glutamine via oxidative decarboxylation versus reductive carboxylation of glutamine. **B** Malate m+3 abundance, derived from U-^13^C- glutamine (n=9 separate wells were sampled per genotype). **C** malate m+3 : malate m+2 ratio, derived from U-^13^C-glutamine (n= 9 separate wells were sampled per genotype). **D** AS m+3: AS m+2 ratio, derived from U-^13^C-glutamine (n= 9 separate wells were sampled per genotype). All P-values were determined using a one-way ANOVA test with Sidak multiple comparisons test. Error bars indicate SD. Measure of centrality is mean.

**Extended Data Figure Figure 6.**
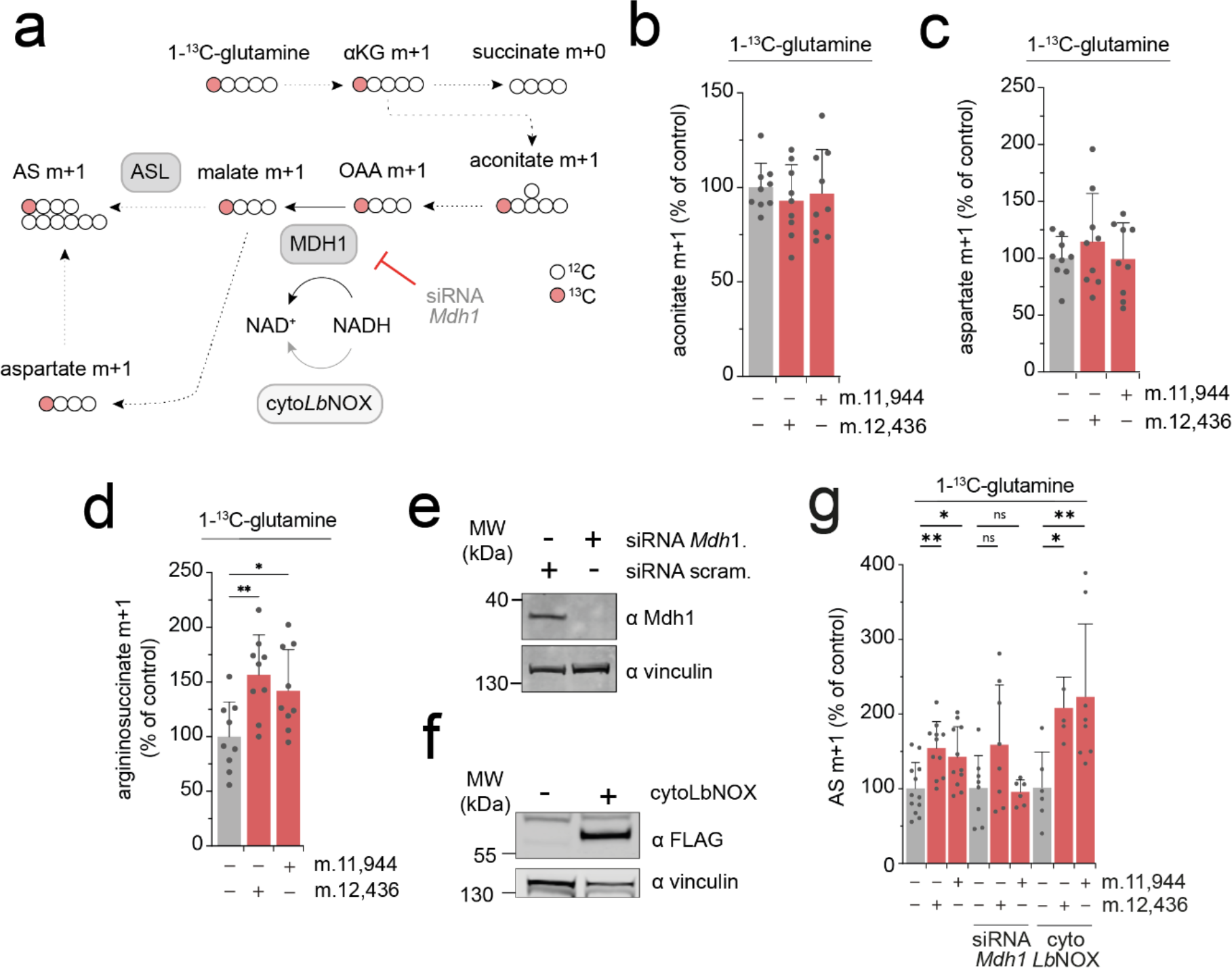
**Increased malate abundance occurs at the level of MDH1 but is not directly due to cytosolic redox potential**. **A** Labeling fate of ^13^C derived from 1-^13^C- glutamine which exclusively labels metabolites derived from the reductive carboxylation of glutamine. **B** Aconitate m+1 abundance, derived from 1-^13^C- glutamine (n= 9 separate wells were sampled per genotype). **C** Aspartate m+1 abundance, derived from 1-^13^C-glutamine (n= 9 separate wells were sampled per genotype). **D** AS m+1 abundance, derived from 1-^13^C-glutamine (n= 9 separate wells were sampled per genotype). **E** Immunoblot of siRNA mediated depletion of *Mdh1*. Representative image shown. **F** Immunoblot of cytoLbNOX expression 36hrs post-sort, detected using αFLAG. Representative image shown. **G** AS m+1 abundance, derived from 1-^13^C-glutamine with indicated treatment (n = 6-12 separate wells were sampled per genotype per condition). All P-values were determined using a one-way ANOVA test with Sidak multiple comparisons test. Error bars indicate SD. Measure of centrality is mean.

**Extended Data Figure Figure 7.**
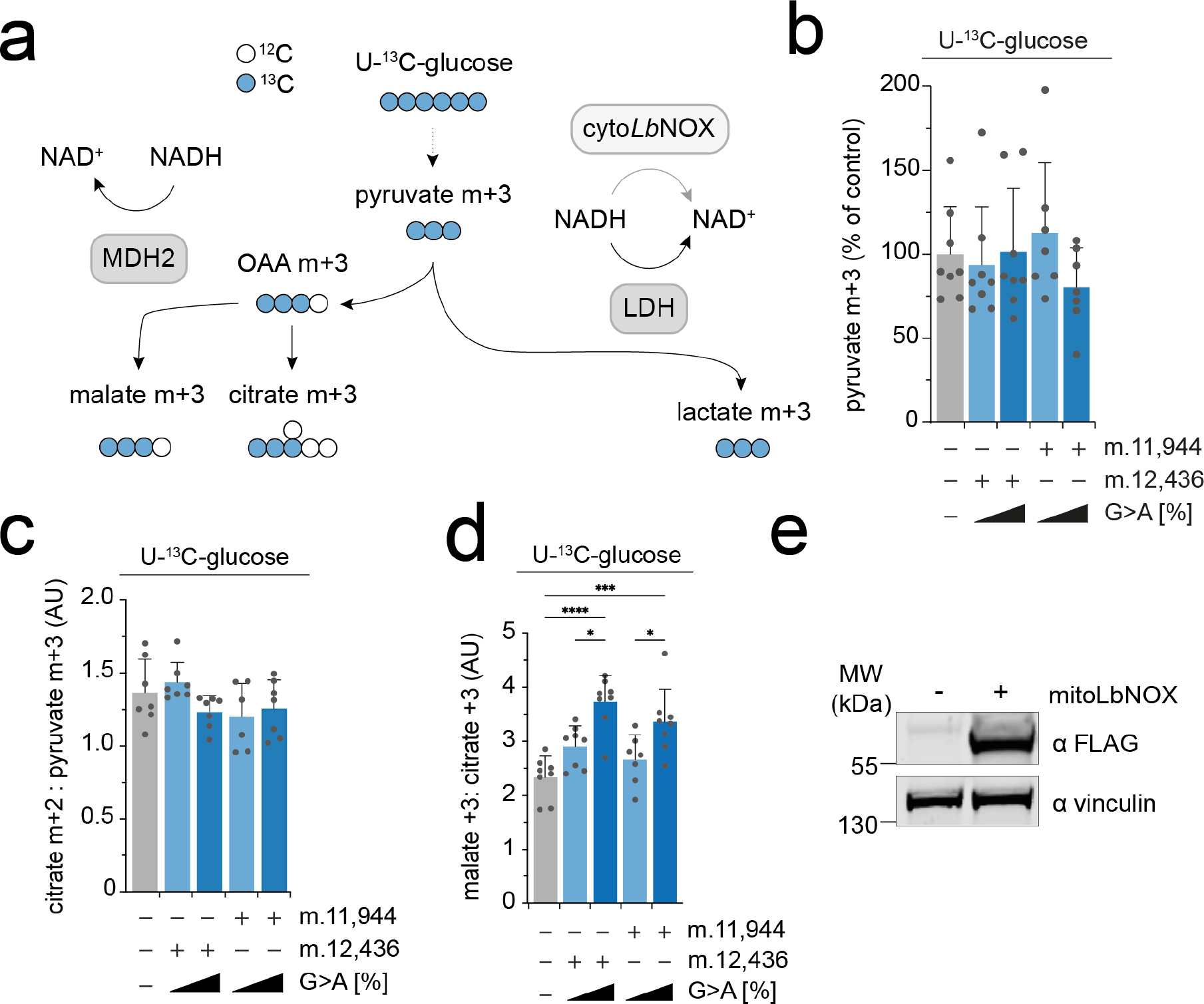
**Increased malate abundance in mutant cells is partially due to MDH2 reversal**. **A** Labeling fate of ^13^C derived from U-^13^C-glucose. **B** Pyruvate m+3 abundance, derived from U-^13^C-glucose (n = 7-8 separate wells were sampled per genotype). **C** Citrate m+2 : pyruvate m+3 ratio, derived from U-^13^C-glucose (n = 6-7 separate wells were sampled per genotype). **D** Malate m+3 : citrate m+3 ratio, derived from U-^13^C-glucose (n = 7-8 separate wells were sampled per genotype). **E** Immunoblot of mitoLbNOX expression 36hrs post-transfection, detected using αFLAG. Representative image shown. All P-values were determined using a one-way ANOVA test with Sidak multiple comparisons test. Error bars indicate SD. Measure of centrality is mean.

**Extended Data Figure Figure 8.**
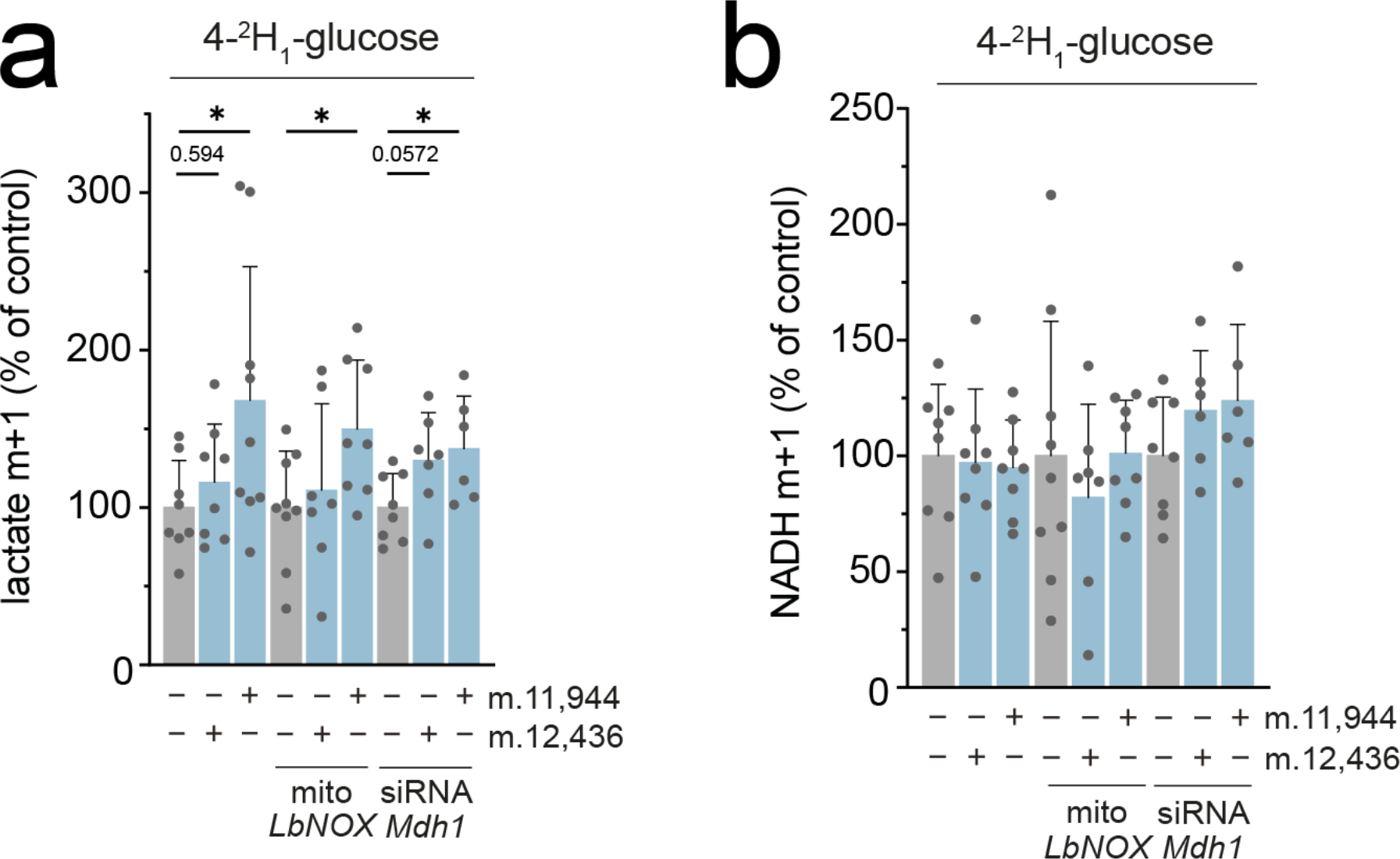
**4-^2^H1-glucose tracing demonstrates that shuttling of electrons between MDH1 and GAPDH drives aerobic glycolysis**. **A** Lactate m+1 abundance, derived from 4-^2^H1-glucose with indicated treatment (n = 7-9 separate wells were sampled per genotype per condition). **B** NADH m+1 abundance, derived from 4-^2^H1-glucose with indicated treatment (n = 6-8 separate wells were sampled per genotype per condition). All P-values were determined using a one-way ANOVA test with Sidak multiple comparisons test. Error bars indicate SD. Measure of centrality is mean.

**Extended Data Figure Figure 9.**
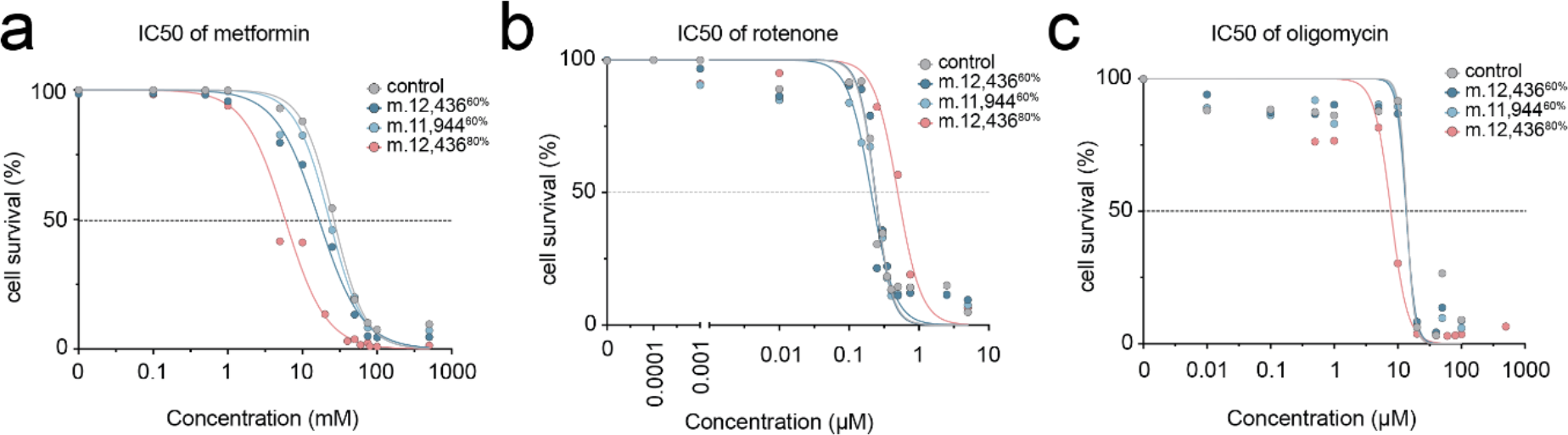
Mutant cells demonstrate a heteroplasmy dose-dependent sensitivity to respiratory chain inhibitors. A. IC50 curve for metformin. IC50 for wild-type = 26.31 ± 1.49mM, for mt.12436^60%^ = 16.60 ± 2.43mM, for mt.12436^80%^ = 5.89 ± 0.71mM and for mt.11944^80%^ = 22.93 ± 0.70mM **B** IC50 curve for rotenone. IC50 for wild-type = 0.236 ± 0.026µM, for mt.12436^60%^ = 0.235 ± 0.035µM, for mt.12436^80%^ = 0.493 ± 0.108µM and for mt.11944^60%^ = 0.205 ± 0.033µM and **C** IC50 curve for oligomycin. IC50 for wild-type = 13.81 ± 3.80µM, for mt.12436^60%^ = 13.52 ± 3.32µM, for mt.12436^80%^ = 7.75 ± 0.56µM and for mt.11944^80%^ = 13.54 ± 3.32µM (n = 4 separate wells per drug concentration per genotype). This was repeated 3 times and a representative result is shown.

**Extended Data Figure Figure 10:**
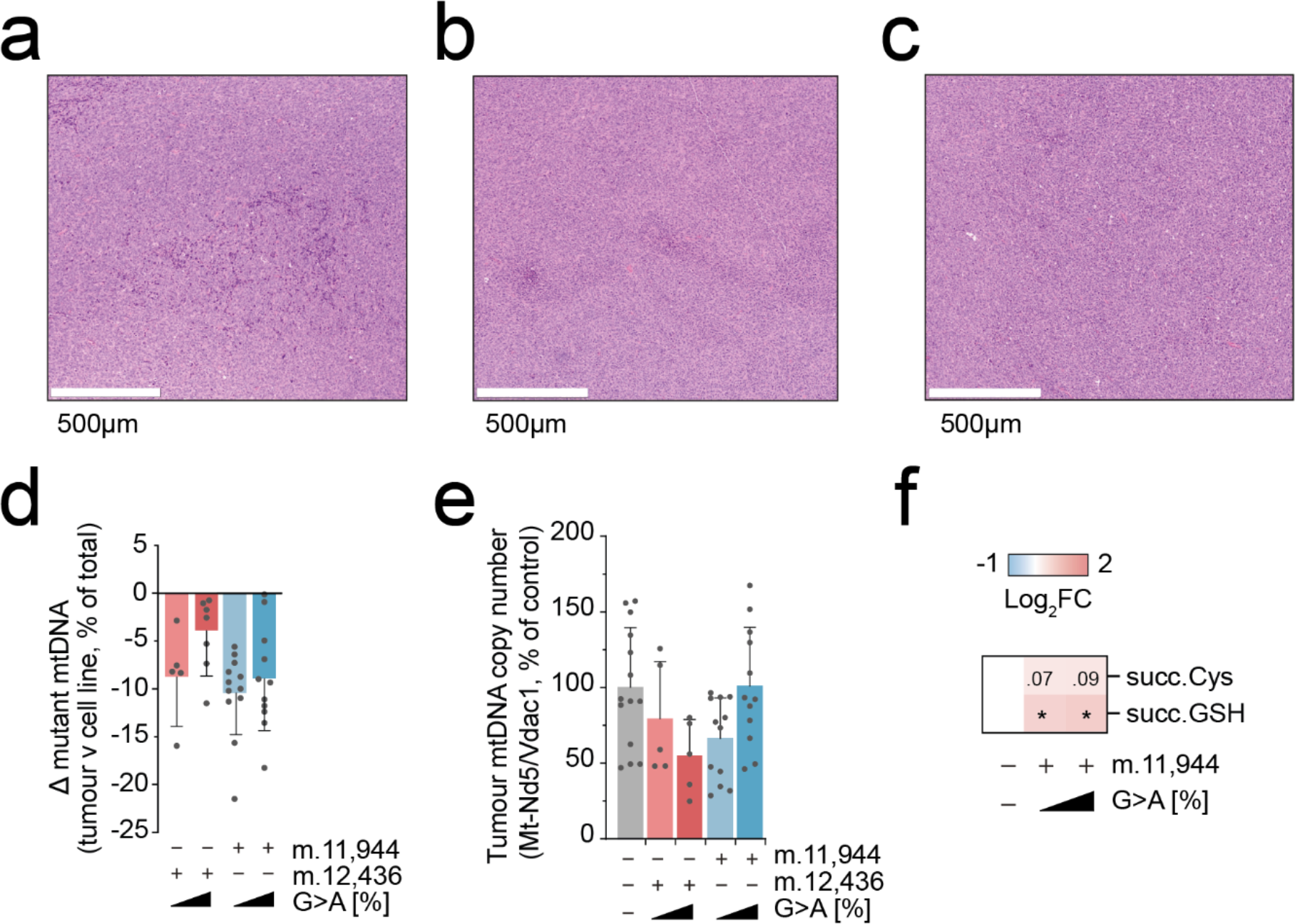
**Allografted B78-D14 lineage tumours do not exhibit macroscopic differences beyond metabolic indicators of disrupted MAS**. Representative H&E sub-section of **A** wild-type, **B** m.12,436^40%^ and **C** m.12,436^60%^ tumours. **D** Change in detected heteroplasmy in bulk tumour samples (n= 5-12 tumours per genotype). **E** Bulk tumour mtDNA copy number (n= 4-13 tumours per genotype). **F** Heatmap of steady-state abundance of metabolically terminal fumarate adducts, succinylcysteine and succinicGSH, demonstrating that metabolic changes observed *in vitro* are preserved *in vivo* (n= 12 tumours per genotype). All P-values were determined using a one-way ANOVA test with Sidak multiple comparisons test. Error bars indicate SD. Measure of centrality is mean.

**Extended Data Figure Figure 11.**
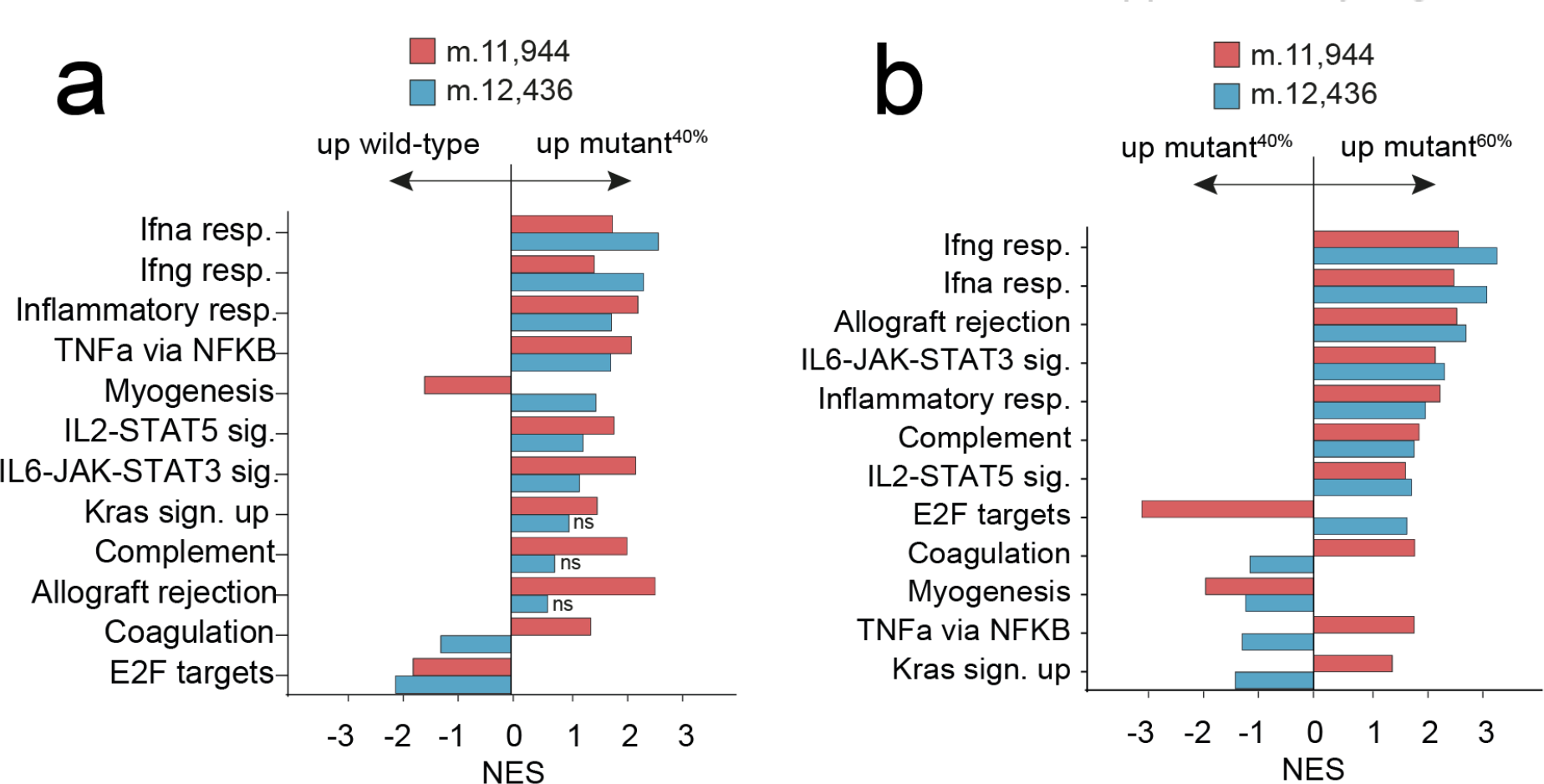
Bulk tumour transcriptional signatures show dose-dependent, heteroplasmy changes in immune-relevant transcriptional phenotypes. GSEA of bulk tumour RNAseq data (n=5-6 tumours per genotype ) showing **A** mutant^40%^ versus wild- type and **B** mutant^60%^ versus mutant^40%^. Only genesets with adj. p-value <0.1 are shown unless otherwise stated. Wilcoxon signed rank test applied.

**Extended Data Figure Figure 12.**
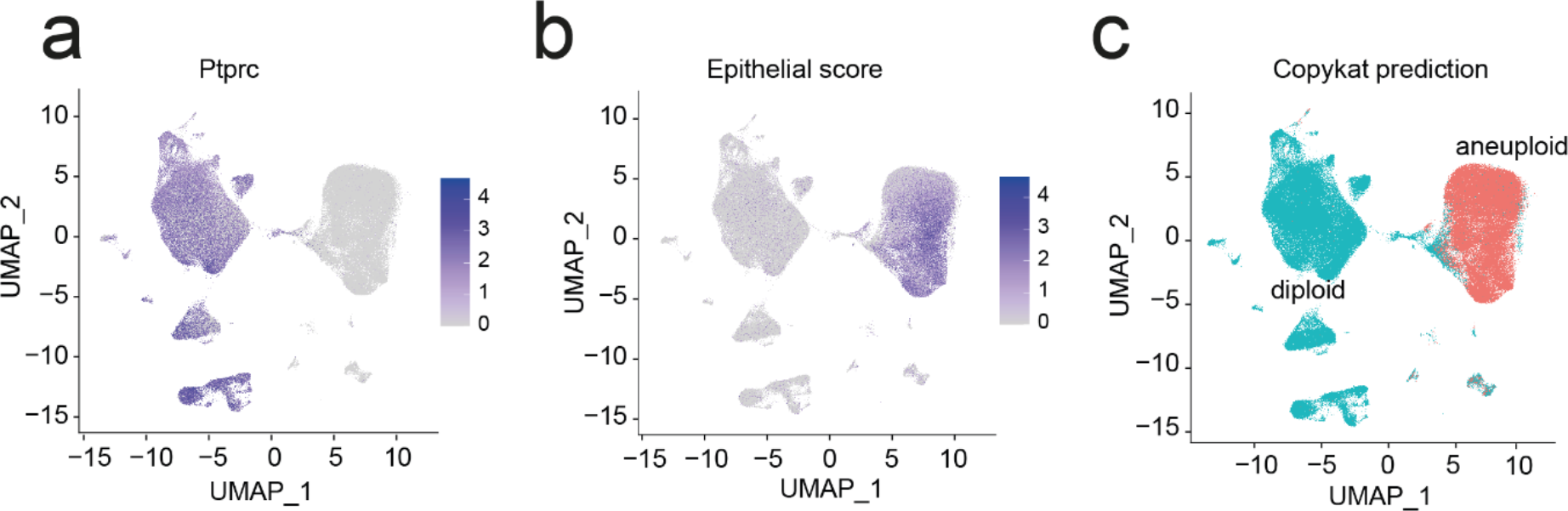
Malignant cells were defined in scRNAseq analysis as aneuploid cells with low or nil Ptprc (CD45) expression and high epithelial score. UMAP indicating **A** Ptprc expression, **B** epithelial score and **C** aneuploidy as determined by copykat prediction. These criteria were employed as the B78 cells lack distinct transcriptional signatures.

**Extended Data Figure Figure 13.**
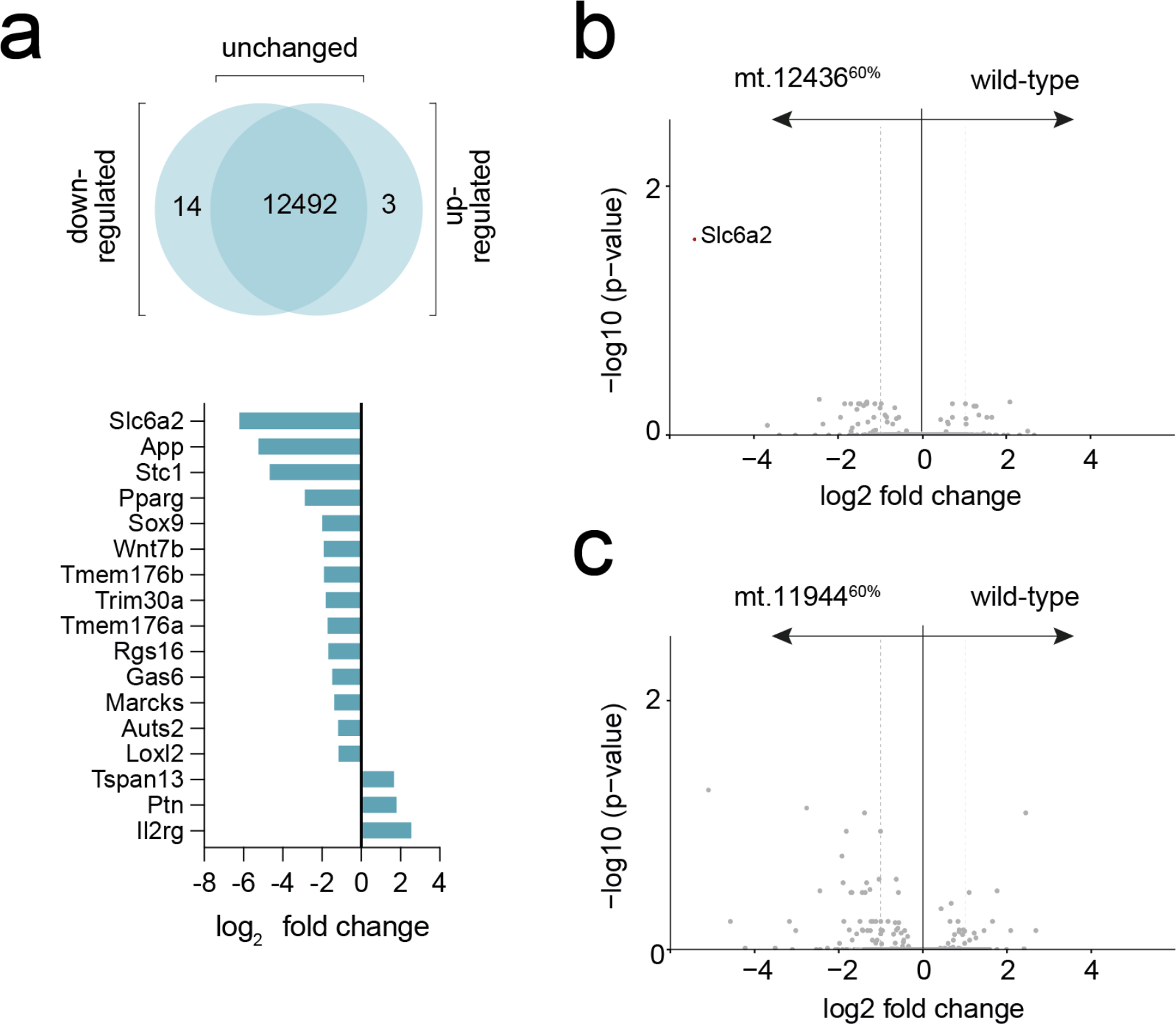
Mutant cells did not have significant changes in transcriptional signatures *in vitro*. **A** Significantly co-regulated transcripts from combined 60% mutant cells versus wild-type (n=12 cell pellets were sampled per genotype). Volcano plot showing differences in gene expression of **A** mt.12436^60%^ cells and **B** mt.11944^60%^ cells versus wild-type. Differences of p < 0.05 and log2 fold change > 1 shown in red (n=12 separate wells were sampled). Wilcoxon signed rank test applied.

**Extended Data Figure Figure 14.**
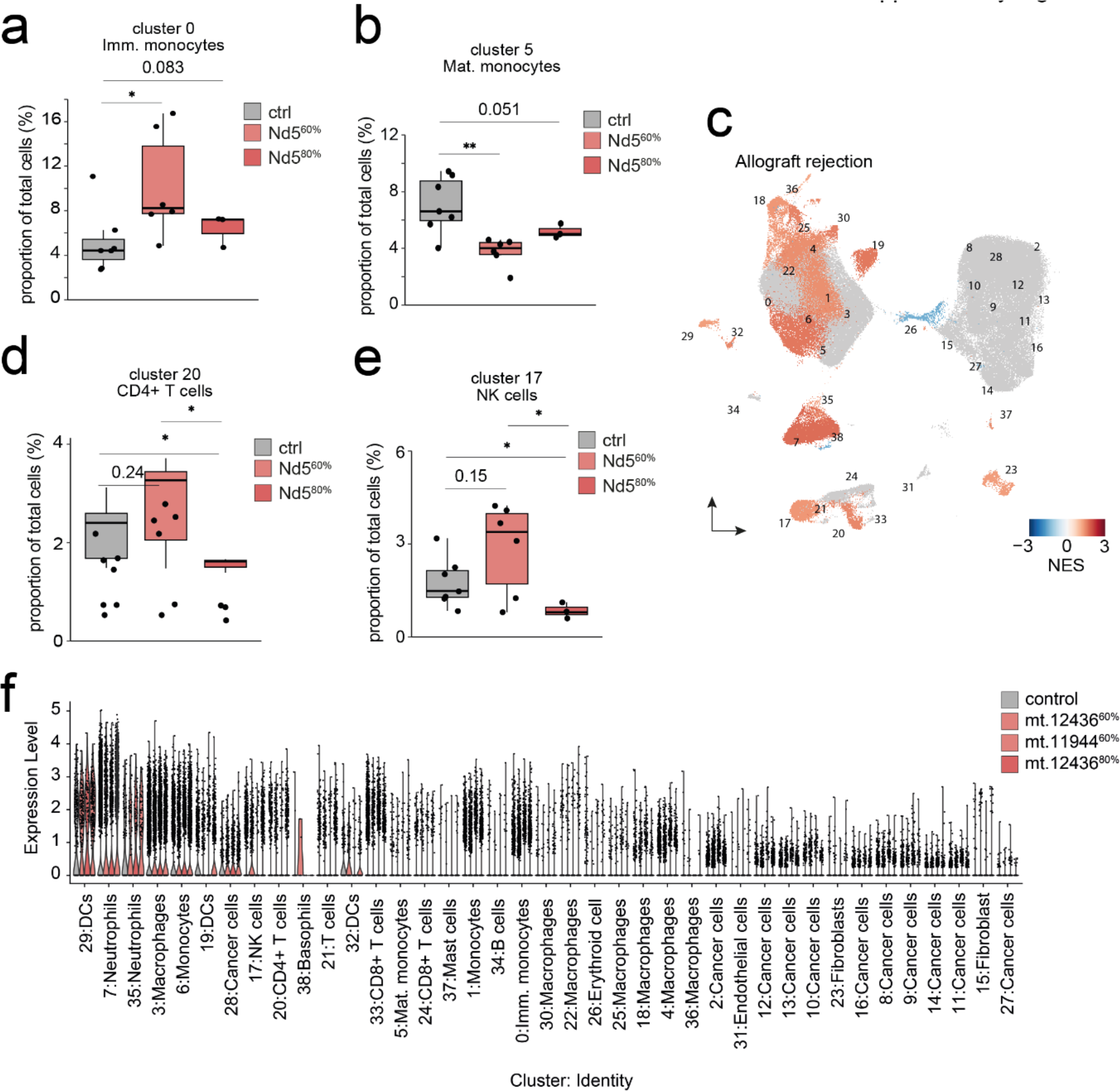
scRNAseq analyses reveal distinct alterations in the tumour immune microenvironment of mtDNA mutant tumours. Proportion of tumour resident: **A** immature monocytes; and **B** CD4+ T-cells relative to the total malignant and non-malignant cells (n = 3-7 tumours per genotype). **C** UMAP coloured by GSEA NES score for allograft rejection geneset. Proportion of tumour resident: **D** CD4+ T cells; and **E** natural killer (NK) cells relative to the total malignant and non-malignant cells (n = 3-7 tumours per genotype). **F** Relative PD-L1 expression within each cell (n = 3- 7 tumours per genotype). One-way ANOVA test with Wilcoxon signed rank test (A) and two-tailed student’s t-test (A-B, D-E) were applied. Error bars indicate SEM. Measure of centrality is mean. Box plots indicate interquartile range (A-B, D-E). NES: normalised expression score. DC, dendritic cell.

**Extended Data Figure Figure 15.**
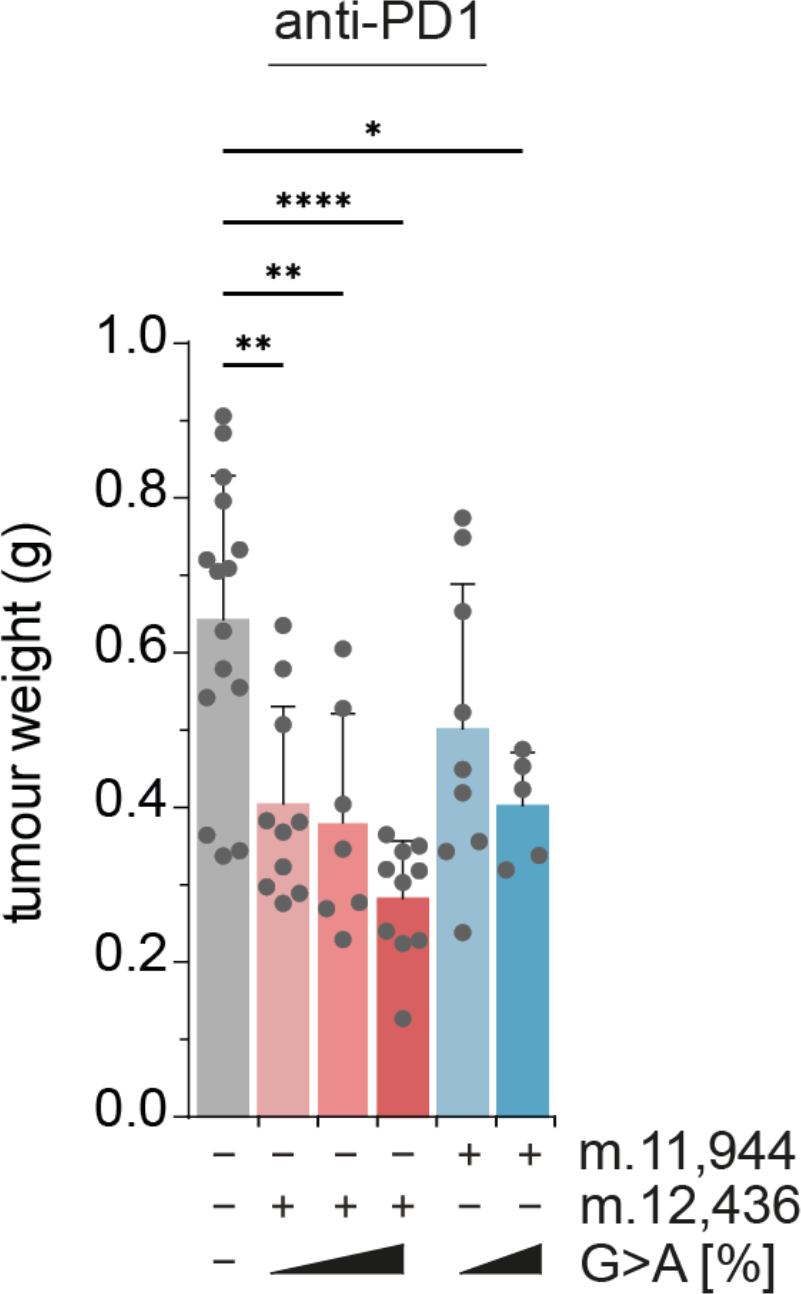
Remodelling of the tumour microenvironment in mutant cells sensitizes tumours to checkpoint blockade. Harvested tumour weight at day 21 (n= 5-15 tumours per genotype). One-way ANOVA test with Sidak multiple comparisons test was applied. Error bars indicate SD. Measure of centrality is mean.

**Extended Data Figure Figure 16.**
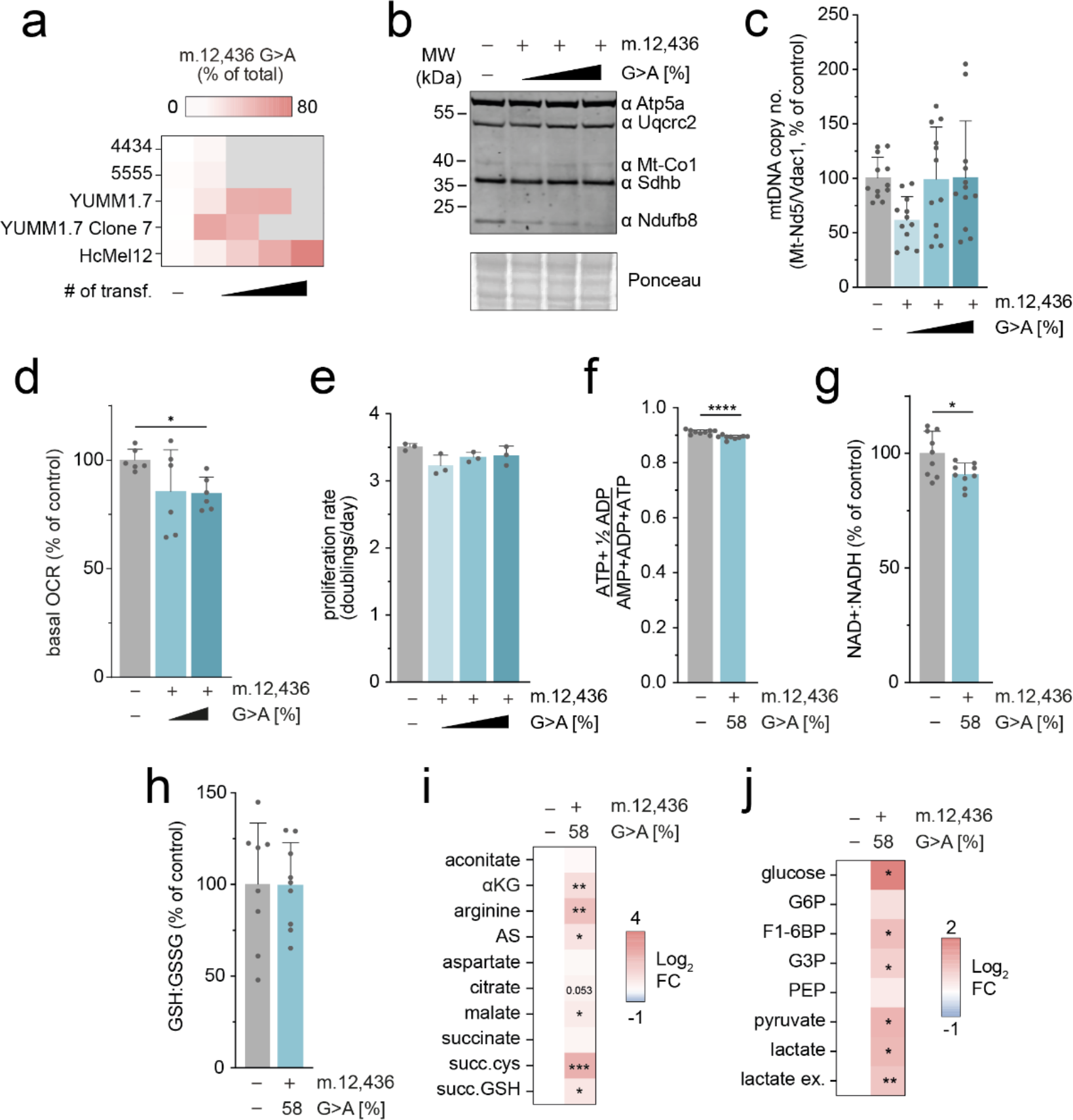
HcMel12 mutant cells recapitulate the cellular and metabolic phenotypes observed in B78-D14 cells. A. Heteroplasmy changes upon subsequent transfection of melanoma cell lines (n= 3 separate cell pellets per genotype). **B** Immunoblot of indicative respiratory chain subunits. Representative result is shown. **C** mtDNA copy number (n= 12 separate wells per genotype). **D** Basal oxygen consumption rate (OCR) (n = 6 measurements (12 wells per measurement) per genotype). **E** Proliferation rate of cell lines in permissive growth media (n = 3 separate wells per genotype) **F** Energy (adenylate) charge state (n = 9 separate wells per genotype). **G** NAD+:NADH ratio (n= 9 separate wells per genotype). **H** GSH : GSSG ratio (n= 8-9 separate wells per genotype). **I** Heatmap of unlabelled steady-state abundance of select mitochondrial metabolites, arginine, argininosuccinate (AS) and terminal fumarate adducts succinylcysteine (succ. Cys) and succinicGSH (succ.GSH) (n= 9 separate wells per genotype). **J** Heatmap of unlabelled steady-state metabolite abundances for select intracellular glycolytic intermediates and extracellular lactate (ex. lactate) (n= 9 separate wells per genotype). P-values were determined using a one-way ANOVA test with (C-D) Sidak multiple comparisons test, Fisher’s LSD Test (E)or (F-J) a one-tailed student’s t-test. Error bars indicate SD. Measure of centrality is mean.

**Extended Data Figure Figure 17.**
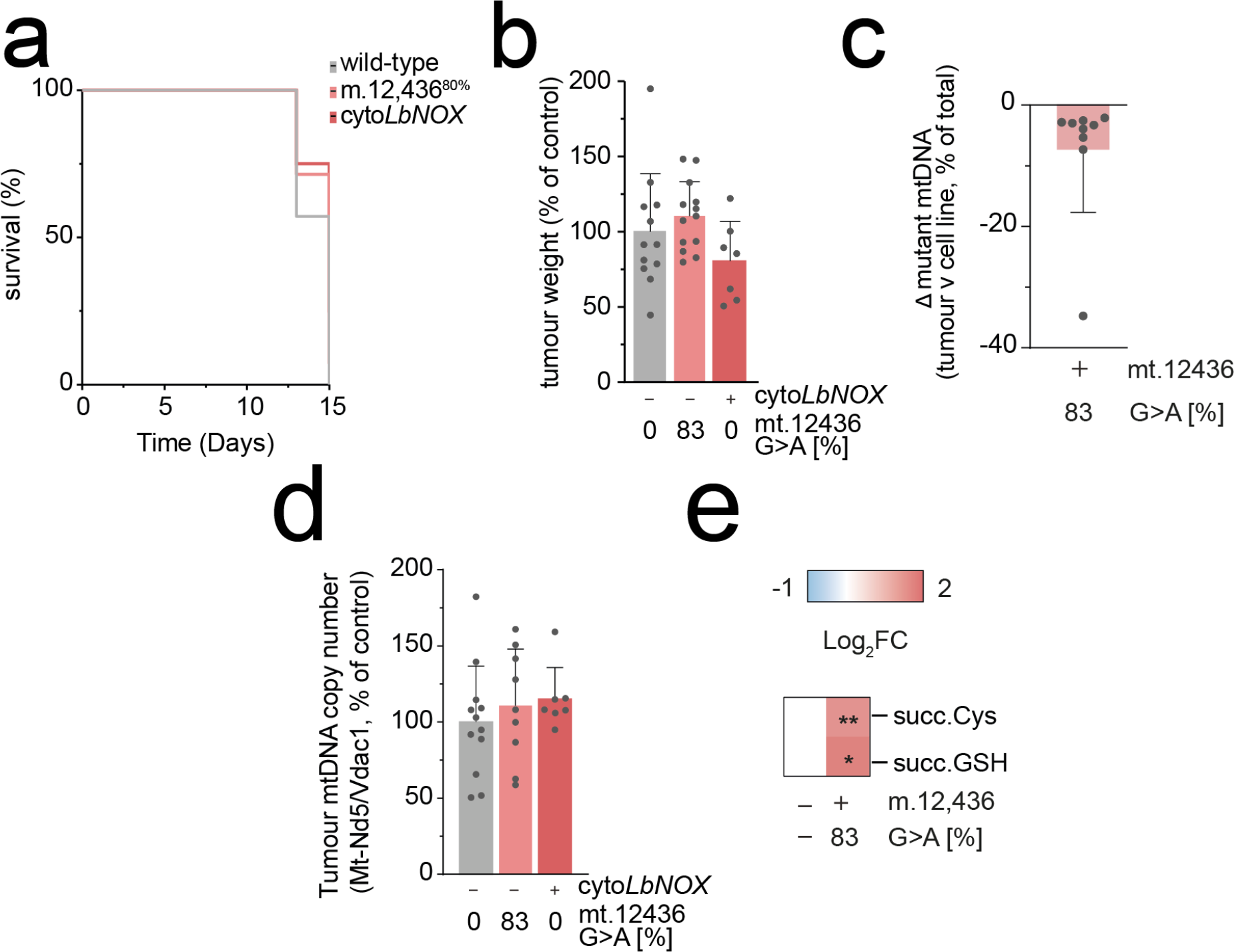
Untreated Hcmel12 lineage tumours recapitulate B78-D14 lineage. A. Survival of C57/BL6 mice subcutaneously injected with indicated cells (n = 9-10 animals per genotype). **B** Tumour weight at endpoint (n = 9-10 tumours per genotype). **C** Change in detected heteroplasmy in bulk tumour samples (n= 9 tumours per genotype). **D** Bulk tumour mtDNA copy number (n= 9 tumours per genotype). **E** Heatmap of steady-state abundance of metabolic terminal fumarate adducts, succinylcysteine and succinicGSH, demonstrating that metabolic changes observed in B78 mutant tumours are preserved *in vivo* (n= 9 tumours per genotype). P-values were determined using a one-way ANOVA test with (B,D) Sidak multiple comparisons test or student’s one-tailed t-test (E). Error bars indicate SD. Measure of centrality is mean.

**Extended Data Figure Figure 18.**
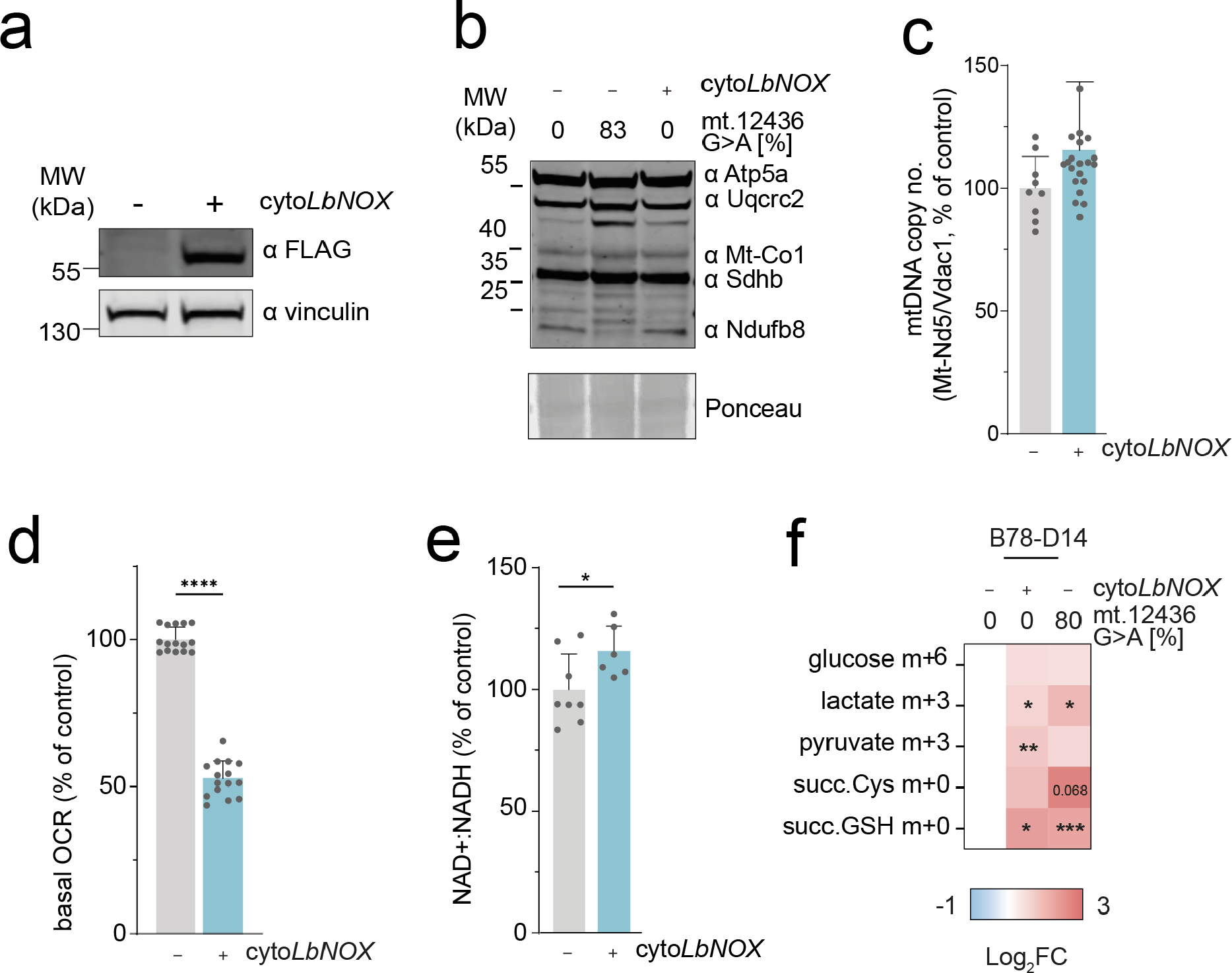
Constitutive expression of cyto*Lb*NOX phenocopies metabolic changes observed in *mt-Nd5* mutant cells. A. Immunoblot of cyto*Lb*NOX expression in clonal population, detected using αFLAG. Representative image shown. **B**. Immunoblot of indicative respiratory chain subunits. Representative result is shown. **C.** mtDNA copy number (n= 9 separate wells per genotype). **D** Basal oxygen consumption rate (OCR) (n = 9-15 measurements (6 wells per measurement) per genotype) A significant decrease is observed in HcMel12 cyto*Lb*NOX, akin to the decrease in basal OCR measured in m.12,436^80%^ cells. **E.** NAD+:NADH ratio (n= 11-12 separate wells per genotype ). **F.** Heatmap of metabolite abundance of glucose m+3, lactate m+3, pyruvate m+3, and terminal fumarate adducts succinylcysteine (succ. Cys) and succinicGSH (succ.GSH) in U- ^13^C-glucose labelling of B78 cells. B78 wild-type cells were transiently transfected with cyto*Lb*NOX and metabolites were extracted 3 days post-sort. A significant increase in lactate abundance was observed in cyto*Lb*NOX-expressing cells mimicking that observed in m.12,436^80%^ cells. (n= 9-13 separate wells per genotype). All P-values were determined using a one-paired student’s t-test. Error bars indicate SD. Measure of centrality is mean.

## Extended Data Methods

### mtDNA sequencing

Cellular DNA was amplified to create two ∼8kbp overlapping mtDNA products using PrimeStar GXL DNA Polymerase (Takara Bio) as per the manufacturer’s instructions.

Primers

Forward 1: 5’-ACTGATATTACTATCCCTAGGAGG-3’

Reverse 1: 5’-TTTGAGTAGAACCCTGTTAGG-3’

Forward 2: 5’-GGCCTGATAATAGTGACGC-3’

Reverse 2: 5’-GGTTGGGTTTAGTTTTTGTTTGG-3’

Resulting amplicons were sequenced using Illumina Nextera kit (150 cycle, paired-end). To determine the percentage of non-target C mutations in mtDNA, we first identified all C/G nucleotides with adequate sequencing coverage (>1000X) in both the reference and experimental sample. Then, for each of the 4 experimental samples, we identified positions for which sequencing reads in the experimental sample corresponded to G>A/C>T mutations. We further filtered the resulting list of mutations to retain only those with a heteroplasmy over 2%, and removed mutations that were also present in control samples. Finally, the non-target percentage was calculated as the fraction of total possible C/G positions that were mutated.

### Sample preparation for MS analysis

Cells were lysed in a buffer containing 4% SDS in 100 mM Tris-HCl pH 7.5 and 55 mM iodoacetamide. Samples were then prepared as previously described in ^23^ with minor modifications. Alkylated proteins were digested first with Endoproteinase Lys-C (1:33 enzyme:lysate) for 1hr, followed by an overnight digestion with trypsin (1:33 enzyme:lysate). Digested peptides from each experimental condition and a pool sample were differentially labelled using TMT16-plex reagent (Thermo Scientific) as per the manufacturer’s instructions. Fully labelled samples were mixed in equal amount and desalted using 100 mg Sep Pak C18 reverse phase solid-phase extraction cartridges (Waters). TMT-labelled peptides were fractionated using high pH reverse phase chromatography on a C18 column (150 × 2.1 mm i.d. - Kinetex EVO (5 μm, 100 Å)) on a HPLC system (LC 1260 Infinity II, Agilent). A two-step gradient was applied, 1% to 28% B (80% acetonitrile) over 42 min, then from 28% to 46% B over 13 min to obtain a total of 21 fractions for MS analysis

### UHPLC-MS/MS analysis

Peptides were separated by nanoscale C18 reverse-phase liquid chromatography using an EASY-nLC II 1200 (Thermo Scientific) coupled to an Orbitrap Fusion Lumos mass spectrometer (Thermo Scientific). Elution was carried out using a binary gradient with buffer A (water) and B (80% acetonitrile), both containing 0.1% formic acid. Samples were loaded with 6 µl of buffer A into a 50 cm fused silica emitter (New Objective) packed in-house with ReproSil-Pur C18-AQ, 1.9 μm resin (Dr Maisch GmbH). Packed emitter was kept at 50 °C by means of a column oven (Sonation) integrated into the nanoelectrospray ion source (Thermo Scientific). Peptides were eluted at a flow rate of 300 nl/min using different gradients optimised for three sets of fractions: 1–7, 8–15, and 16–21^23^. Each fraction was acquired for a duration of 185 minutes. Eluting peptides were electrosprayed into the mass spectrometer using a nanoelectrospray ion source (Thermo Scientific). An Active Background Ion Reduction Device (ESI Source Solutions) was used to decrease air contaminants signal level. The Xcalibur software (Thermo Scientific) was used for data acquisition. A full scan over mass range of 350–1400 m/z was acquired at 60,000 resolution at 200 m/z, with a target value of 500,000 ions for a maximum injection time of 50 ms. Higher energy collisional dissociation fragmentation was performed on most intense ions during 3 sec cycle time, for a maximum injection time of 120 ms, or a target value of 100,000 ions. Peptide fragments were analysed in the Orbitrap at 50,000 resolution.

### Proteomics Data Analysis

The MS Raw data were processed with MaxQuant software^24^ v.1.6.1.4 and searched with Andromeda search engine^25^, querying SwissProt^26^ *Mus musculus* (25,198 entries). First and main searches were performed with precursor mass tolerances of 20 ppm and 4.5 ppm, respectively, and MS/MS tolerance of 20 ppm. The minimum peptide length was set to six amino acids and specificity for trypsin cleavage was required, allowing up to two missed cleavage sites. MaxQuant was set to quantify on “Reporter ion MS2”, and TMT16plex was set as the Isobaric label. Interference between TMT channels was corrected by MaxQuant using the correction factors provided by the manufacturer. The “Filter by PIF” option was activated and a “Reporter ion tolerance” of 0.003 Da was used. Modification by iodoacetamide on cysteine residues (carbamidomethylation) was specified as variable, as well as methionine oxidation and N-terminal acetylation modifications. The peptide, protein, and site false discovery rate (FDR) was set to 1 %. The MaxQuant output ProteinGroup.txt file was used for protein quantification analysis with Perseus software^27^ version 1.6.13.0. The datasets were filtered to remove potential contaminant and reverse peptides that match the decoy database, and proteins only identified by site. Only proteins with at least one unique peptide and quantified in all replicates in at least one experimental group were used for analysis. Missing values were added separately for each column. The TMT corrected intensities of proteins were normalised first by the median of all intensities measured in each replicate, and then by using LIMMA plugin^28^ in Perseus. Significantly regulated proteins between two groups were selected using a permutation-based Student’s t-test with FDR set at 1%.

### Mitochondrial membrane potential and pH gradient

Membrane potential and pH gradient were measured using multi-wavelength spectroscopy as described in ^29–30^ . Briefly, cultured cells were disassociated by gentle tapping and then spun down and resuspended at a density of 1×10^7^ cells/mL in FluroBrite supplemented with 2 mM glutamine in a temperature-controlled chamber. Changes in mitochondrial cytochrome oxidation states were then measured with multi-wavelength spectroscopy. The baseline oxidation state was measured by back-calculation using anoxia to fully reduce the cytochromes, and a combination of 4μM FCCP and 1μM rotenone to fully oxidize the cytochromes. The membrane potential was then calculated from the redox poise of the b-hemes of the *bc1* complex and the pH gradient measured from the turnover rate and redox span of the *bc1* complex using a model of turnover^30^.

### Mitochondrial NADH oxidation state

Changes in NAD(P)H fluorescence were measured simultaneously with mitochondrial membrane potential using 365nm excitation. The resultant emission spectrum was then measured with multi-wavelength spectroscopy^29^. The baseline oxidation state of the mitochondrial NADH pool was back calculated using anoxia to fully reduce, and 4 μM FCCP to fully oxidize the mitochondrial NADH pool, respectively, assuming the cytosolic NADH pool and NADPH pools did not change with these interventions and short time period.

### H&E Staining

Haematoxylin and Eosin (H&E) staining and slide scanning was performed as described in ^31^.

### scRNAseq Methodology

1- Preprocessing of single-cell RNA transcriptomics data, batch effect correction, and clustering

CellRanger (v.7.0.1) was used to map the reads in the FASTQ files to the mouse reference genome (GRCm39)^32^. Seurat (v.4.2.0) package in R (v.4.2.1) was used to handle the pre-processed gene counts matrix generated by cellRanger^33^. As an initial quality control step, cells with fewer than 200 genes as well as genes expressed in less than 3 cells were filtered out. Cells with >5% mitochondrial counts, UMI counts > 37000, and gene counts < 500 were then filtered out. The filtered gene counts matrix (31647 genes and 127356 cells) was normalized using the NormalizeData function using the log(Normalization) method and scale.factor to 10000. The FindVariableFeatures function was used to identify 2000 highly variable genes for principal component analysis. The first 50 principal components were selected for downstream analysis. RunHarmony function from harmony package (v.0.1.0 ) with default parameters was used to correct batch effects^34^. The RunUMAP function with the reduction from “harmony” was used to generate UMAPs for cluster analysis. FindClusters function was used with the resolution parameters set to 1.6.

2- Epithelial score

Average gene expression from cytokeratins, Epcan, and Sfn were used to calculate epithelial score.

3- Single-cell copy number estimation

CopyKat (v.1.1.0) was used to estimate the copy number status of each cell^14^. Parameters were set as ngene.chr=5, win.size=25, KS.cut=0.1, genome=”mm10” and cells annotated as T cells or NK cells in the UMAP as diploid reference cells.

4- Identification of differential expressed marker genes

Top differentially expressed genes in each cluster were identified using the FindAllMarkers function in the Seurat R package. Parameters for expression difference were set to be at least 1.25 times of fold changes (logfc.threshold = 1.25) and adjusted p-value < 0.05 with gene expression detected in at least 10% of cells in each cluster (min.pct = 0.1). The top 20 highly differentially expressed genes in each cluster ranked by average fold change were defined as marker genes.

5- Pathway enrichment analysis of single-cell transcriptomics data

For cells in each identified cluster in the UMAP, the wilcoxauc function from presto R package ( version 1.0.0) was used to conduct wilcox rank-sum test to obtain the fold change and p-value for all genes between cells in the high heteroplasmy group for both mutations and control group^35^. The genes were ranked in decreasing order according to the formula sign(log2FC) * (-log10(p-value) ). This ranked gene list, and mouse hallmark pathways (mh.all.v2002.1.Mm.symbols.gmt) from the MSigDB database were used as inputs for gene set enrichment analysis using the fgsea function from fgsea R package (v.1.22.0) with parameters of eps=0, minSize=5, maxSize =500^36^.

